# TSPAN3 defines a distinct intracellular trafficking route to secretory multivesicular bodies

**DOI:** 10.64898/2026.08.26.747209

**Authors:** Jelle van den Bor, Misko Bobeldijk, Christopher A. Zala, M. Cristina Trueba Sanchez, Clara Lalo, Bárbara Adem, Jochanan A. Maaijen, Emma M. A. Bundock, Nikki A. Weijers, Elly Soltani, Cecilia de Heus, Pascal W.T.C. Jansen, Wenyi Zheng, Samir El Andaloussi, Nalan Liv, Annemiek van Spriel, Kelly Stecker, Ihor SmaI, Guido van Mierlo, Frederik J. Verweij

## Abstract

Extracellular vesicles (EVs) comprise molecularly diverse populations generated through multiple membrane-trafficking pathways, yet the intracellular basis of this heterogeneity remains poorly understood. Here, we identify the EV-associated tetraspanin TSPAN3 as a marker of a secretory multivesicular body (MVB) population that is molecularly and functionally distinct from canonical CD63-positive compartments. Using endogenous genome editing, live-cell and super-resolution microscopy, electron microscopy, quantitative EV secretion assays, and complementary proteomic approaches, we show that TSPAN3 localizes to fusion-competent MVBs but exhibits limited overlap with CD63 during secretion. Unlike CD63, which extensively traffics through the plasma membrane and depends on YXXΦ-mediated endocytic retrieval, TSPAN3 reaches secretory MVBs predominantly through an intracellular trafficking route that relies on a dileucine-containing sorting region. Orthogonal proximity-labeling and affinity-purification proteomics revealed that TSPAN3-positive compartments are associated with a selective LC3/ATG8-related membrane network, including GABARAPL2 and proteins involved in endosomal membrane remodeling and fusion. Perturbation of residues required for this association impaired localization to LC3-positive compartments and reduced secretory MVB fusion. Consistent with these findings, pharmacological disruption of autophagy-and endolysosomal-associated pathways differentially altered TSPAN3-positive EV secretion. Finally, proximity-labeled EV proteomics demonstrated that TSPAN3-associated EVs possess cargo signatures distinct from CD63-associated EVs, with greater representation of endosomal and endolysosomal proteins suggesting that tetraspanin-associated membrane nanodomains retain molecular signatures consistent with their intracellular trafficking history. Together, our findings identify TSPAN3 as a marker of a previously unrecognized secretory MVB population distinguished by its intracellular trafficking, molecular interactions, and EV composition, supporting a model in which distinct tetraspanin-organized membrane nanodomains are associated with different intracellular trafficking routes and molecularly distinct EV populations.

## Introduction

Extracellular vesicles (EVs) comprise a highly heterogeneous population of membrane-enclosed particles that differ in size, molecular composition and cellular origin. This heterogeneity poses a major challenge for the identification and isolation of biologically distinct EV populations. Tetraspanins, particularly CD9, CD63 and CD81, are among the most widely used EV-associated proteins and are routinely employed for EV characterization and affinity-based isolation. However, increasing single-vesicle analyses have demonstrated that these canonical tetraspanins are not uniformly distributed across EVs. Distinct CD9-, CD63-and CD81-positive EV populations can be resolved within individual samples, while a substantial fraction of EVs lacks detectable levels of all three markers^1–3^. Indeed, multiplexed single-EV profiling recently showed that ∼40% of EVs lacked detectable CD9, CD63 and CD81, whereas only a small minority expressed all three^3^. These findings indicate that commonly used tetraspanin markers fail to capture a substantial fraction of EVs, highlighting the need for additional markers to resolve previously unrecognized EV populations. Given the established role of tetraspanins as organizers of membrane protein nanodomains, other, less-studied members of the tetraspanin family may likewise define molecularly distinct EV populations.

Consistent with this, a large-scale screen of 244 candidate EV-sorting proteins identified TSPAN2 and TSPAN3 as particularly efficient EV-associated scaffold proteins across multiple producer cell types, with both outperforming CD63 in several reporter-based assays^4^. TSPAN2-and TSPAN3-engineered EVs also showed limited representation of the canonical tetraspanins CD9, CD63 and CD81 and displayed proteomic profiles distinct from CD63-engineered EVs, suggesting that these tetraspanins may associate with molecularly distinct EV populations^4^. More recently, systematic definition of reference small-EV proteomes identified TSPAN3 among the top 100 sEV-associated proteins, further supporting its broad association with extracellular vesicles^5^. These observations suggest that TSPAN3 may not merely represent an additional EV marker, but may instead identify a molecularly distinct secretory pathway responsible for generating a subset of EVs. However, the mechanisms governing TSPAN3 trafficking to EV-producing compartments and its incorporation into EVs remain largely unknown. In particular, it is unclear whether TSPAN3 localizes to secretory multivesicular bodies (MVBs) and which trafficking machinery directs its incorporation into EVs.

Understanding how TSPAN3 becomes incorporated into EVs requires knowledge of how EV-associated tetraspanins traffic through the endolysosomal system. CD63, one of the best-characterized EV-associated tetraspanins, dynamically traffics between the plasma membrane and endosomal compartments and is efficiently retrieved from the cell surface through adaptor protein-dependent sorting mediated by its cytosolic YXXΦ motif^6,7^. Following endocytosis, CD63 accumulates in late endosomes and multivesicular bodies (MVBs), where inward budding of the limiting membrane generates intraluminal vesicles (ILVs). MVBs can subsequently fuse with lysosomes for degradation or with the plasma membrane to release ILVs as exosomes^8,9^. However, depending on the cell type and EV biogenesis pathway, CD63 can also be incorporated efficiently into EVs by directly budding from the plasma membrane, indicating that its presence is not restricted to endosome-derived vesicles^10^. Thus, the intracellular trafficking routes followed by individual tetraspanins may influence whether they enter the endosomal system, the specific MVB populations into which they are sorted, and ultimately their incorporation into secreted EVs. Whether less-characterized EV-associated tetraspanins such as TSPAN2 and TSPAN3 use the same trafficking route and co-reside in the same compartments as CD63 remains unknown.

Increasing evidence indicates that MVBs themselves constitute heterogeneous populations with distinct molecular identities and cellular fates. Secretory competence appears to be acquired by specific MVB subsets during endosomal maturation; for example, plasma membrane-fusing CD63-positive MVBs have been identified as a Rab27-positive pre-endolysosomal population that is molecularly distinct from the majority of intracellular CD63-positive compartments^8^. Other secretory MVB populations are distinguished by their Rab GTPase composition, lipid identity, and molecular machinery, while specific trafficking regulators can bias MVBs toward either plasma membrane fusion or lysosomal degradation. Thus, EV heterogeneity may arise not only through differential cargo sorting into ILVs, but also through the selective maturation and fate of distinct parental MVBs. However, the molecular features that specify secretory MVB identity, and how individual EV-associated tetraspanins become linked to particular secretory compartments, remain incompletely understood.

Despite the widespread use of tetraspanins as EV markers, it remains unclear how the intracellular trafficking of individual tetraspanins relates to the molecular identity and secretory behavior of the compartments they occupy. Here, we investigated TSPAN3 as a poorly characterized EV-associated tetraspanin, initially comparing its localization and EV secretion with established EV-associated tetraspanins. Using endogenous tagging, live-cell and TIRF microscopy, ultrastructural analysis, and complementary proximity-labeling and affinity-purification proteomics, we identify TSPAN3 as a marker of a secretory MVB population distinct from canonical CD63-positive compartments. We further show that TSPAN3 and CD63 differ in their intracellular trafficking requirements and uncover an unexpected association between TSPAN3-positive compartments and LC3/ATG8-related machinery. Together, our findings establish TSPAN3 as a molecular handle for resolving heterogeneity within the secretory endosomal system and provide a framework for understanding how intracellular trafficking contributes to EV diversity.

## Methods

### Cell culture and pharmacological treatments

HeLa FlpIN cells were maintained in Dulbecco’s modified Eagle’s medium (DMEM) supplemented with 10% fetal bovine serum (FBS) and 1% penicillin–streptomycin at 37°C in a humidified atmosphere containing 5% CO₂. Cells were routinely tested for mycoplasma contamination.

For pharmacological perturbation experiments, cells were treated for 18 h with Bafilomycin A1 (BafA1; MedChemExpress; 100 µM), SAR405 (MedChemExpress; 1 µM), ATG7-IN-2 (MedChemExpress; 1 µM), dynasore (MedChemExpress; 150 µM), or Pitstop 2 (MedChemExpress; 5 µM). For combined treatments, compounds were added simultaneously unless stated otherwise. Vehicle-control cells were treated with the corresponding concentration of DMSO.

### Generation of endogenous HA-Nluc and HALO knock-in cell lines

Endogenous HA-Nanoluc-tetraspanin knock-in cell lines were generated by in-trans paired nicking (ITPN) CRISPR/Cas9-mediated genome editing as described by Bollen et al., 2022^11^. In brief, Guide RNAs targeting the N-terminus of TSPAN2 (5’-CGCGGAAGCGCCCCATGCTG-3’), TSPAN3 (5’-GGTGATGCCGCACTGGCCCA3-’), CD9 (5’-GGTGCCTCCTTTGACCGGCA-3’), and CD63 (5’-GCAGCCATGGCGGTGGAAGG-3’) were designed using the UCSC Genome Browser^12^. HeLa FlpIN wild-type cells were transfected with a PX462^13^ plasmid containing dCas9 and the specific sgRNA, and a template plasmid containing the HA-NLuc cassette that is flanked by gene-specific 600-base-pair upstream and downstream homology arms^11^. After puromycin selection (2 µg/mL), single-cell clones were seeded into 96-well plates and screened for secreted NLuc activity. Positive clones were isolated and correct integration was validated by Sanger sequencing.

For endogenous fluorescence imaging, we introduced a HALO tag into the first extracellular loop (ECL1) of TSPAN2, TSPAN3, CD9, and CD63. For TSPAN3 and CD63, we generated ECL1-HALO knock-ins using homology-mediated end-joining^14^ (HMEJ), whereas for TSPAN2 and CD9 we used the ITPN strategy as listed above^11^. As described above, guides targeting ECL1 for TSPAN2 (5’-TACTCGGTTCGGAGGTGCCATAA-3’), TSPAN3 (5’-TGTACACATCTTCAAAGAAG-3’), CD9 (5’-TTCTTGCTCGAAGATGCTCT3-’), CD63 (5’-CTGAGTCAGACCATAATCCA3-’). For the HMEJ strategy, HeLa FlpIN wild-type cells were transfected with a PX459^13^ plasmid containing Cas9 and the specific sgRNA, and a template plasmid containing the HALO tag cassette that is flanked by gene-specific 800-base-pair upstream and downstream homology arms^14^. For the ITPN strategy, we transfected cells as described above. Following puromycin selection (2 µg/mL), single-cell clones were isolated by fluorescence-activated cell sorting after labelling with 1 µM fluorescent JFX554 HALO ligand (Janelia Research Campus). Clonal lines were expanded and characterized by fluorescent HALO-ligand labeling and live-cell fluorescence microscopy to assess expression and subcellular localization. We confirmed correct genomic integration by Sanger sequencing.

### Plasmids and site-directed mutagenesis

CD63, CD9, TSPAN2, and TSPAN3 were expressed as pHluorin fusion proteins in the first extracellular loop for visualization of multivesicular body (MVB)–plasma membrane fusion events. Plasmids for CD63 and CD9 with pHluorin in the first extracellular loop were previously published^15^. pHluorin was integrated in the first extracellular loop-of TSPAN2 and TSPAN3 using Gibson assembly. For dual-color imaging of CD63-and TSPAN3-positive compartments, CD63 was fused to pHmScarlet and using in combination with the TSPAN3-pHluorin. The plasmid for CD63-pHmScarlet was previously published^15^.

Tetraspanin trafficking mutants and candidate LC3-interacting region (LIR)-associated mutants were ordered at Twist as partial gene fragments and subcloned into the corresponding TSPAN-pHluorin plasmid using restriction enzyme cloning or Gibson assembly. TSPAN3 mutants included T75A, Y252A, LL246AA, and Y244A. CD63 mutant included Y235A. All constructs were verified by Sanger sequencing (Macrogen) and whole plasmid sequencing (Plasmidsaurus)

### NanoLuc-based extracellular vesicle secretion assay

Nanoluc-based extracellular vesicle secretion assay was performed as previously published^16^. In brief, extracellular vesicle secretion was quantified using HeLa cells expressing endogenous HA-Nluc-tagged CD63, CD9, TSPAN2, or TSPAN3. Cells were seeded at 6,000 cells per well in white 96-well plates (Corning, #3917) and allowed to adhere for 24 h before treatment. Cells were subsequently incubated with the indicated pharmacological compounds for 18 h.

Following treatment, conditioned medium was cleared of detached cells and cellular debris by centrifugation at 300 × *g* for 5 min at room temperature. Subsequently, 50 µL of the cleared conditioned medium was transferred to a fresh white 96-well plate (Greiner Bio-One, #07-000-128) for measurement of extracellular NanoLuc activity. The remaining medium was removed from the cells, which were washed with PBS and lysed in assay buffer (100 mM MES hydrate pH 6.0; 1 mM EDTA; 150 mM KCl; 35 mM Thiourea; 0.5% IGEPAL CA-630) freshly supplemented with 1 mM DTT. NanoLuc activity in the conditioned medium and corresponding cell lysates was measured using CTZ-400a (Nanolight Technology, #340) dissolved in NanoFuel-400a solvent (Nanolight Technology, #397) diluted to 10 µM in assay buffer. Luminescence was recorded using a GloMax plate reader (Turner Biosystems Instrument #9101-002, Promega).

For pharmacological perturbation experiments, secreted and cellular NanoLuc activities were analyzed independently. Secreted NanoLuc activity was normalized to the corresponding vehicle-treated condition and is presented as relative EV secretion. Cellular NanoLuc activity was independently normalized to the corresponding vehicle-treated condition and is presented as relative cellular reporter abundance.

In addition, relative secretion efficiency was calculated by dividing secreted NanoLuc activity by the corresponding cellular NanoLuc activity and normalizing this ratio to the vehicle-treated condition.

### Size-exclusion chromatography

Size-exclusion chromatography (SEC) was performed using qEVoriginal 70 nm columns in combination with an Automatic Fraction Collector (AFC; IZON Science). Conditioned medium from HeLa cells expressing endogenous HA-Nluc-tagged CD63, CD9, TSPAN2, or TSPAN3 was collected and cleared of detached cells and cellular debris by centrifugation at 300 × *g* for 5 min at room temperature.

Prior to sample loading, qEVoriginal 70 nm columns were equilibrated with PBS according to the manufacturer’s recommendations. A total sample volume of 500 µL was loaded onto each column, and fraction collection was performed automatically using the IZON AFC. 26 consecutive fractions of 500 µL were collected in 2 mL Eppendorf tubes.

NanoLuc activity was measured in 50 µL of each SEC fraction using CTZ-400a (Nanolight Technology, #340) dissolved in NanoFuel-400a solvent (Nanolight Technology, #397) and diluted to 10 µM in assay buffer. Luminescence was recorded using a GloMax plate reader (Turner Biosystems Instrument #9101-002, Promega). NanoLuc activity in each fraction was expressed as raw luminescence.

EV-associated fractions were identified based on the established elution profile of the qEVoriginal 70 nm column. The distribution of NanoLuc activity across the SEC fractions was used to determine whether extracellular NanoLuc activity was associated with the EV-enriched rather than the soluble-protein fractions.

### Super-resolution fluorescence microscopy

Super-resolution fluorescence imaging was performed using a Zeiss LSM 980 confocal laser-scanning microscope equipped with Airyscan 2 (Carl Zeiss Microscopy). Images were acquired using a Alpha Plan-APO 100x/1,46 WD=0.10mm, Oil objective in Airyscan SR mode. Fluorophores were excited using the appropriate laser lines, and emission was collected using the Airyscan detector. Acquisition settings were kept constant between experimental conditions within individual experiments when fluorescence intensities or localization patterns were compared.

For imaging of endogenous HALO-tagged tetraspanins, cells were labeled with CA-MaP555 HALO ligand at 1 µM for 1 min. Where indicated, the plasma membrane was visualized using fluorescent wheat germ agglutinin (WGA, biotum, CF640R, #29026) at 1 µg/mL for 10 minutes. Cells were subsequently washed with PBS and fixed with 4% paraformaldehyde containing 4% sucrose.

Airyscan images were processed using the Airyscan processing algorithm in Zeiss ZEN Blue using 3D processing settings. Representative images were prepared using FIJI, with identical image-processing settings applied to samples that were quantitatively compared.

### Live-cell TIRF microscopy

Live-cell total internal reflection fluorescence (TIRF) microscopy was performed using a Nikon Eclipse Ti microscope equipped with a Perfect Focus System (PFS; Nikon). Images were acquired using a Nikon Apo TIRF 100×/1.49 NA oil-immersion objective and a Prime BSI sCMOS camera at a spatial sampling of 14.71 pixels/µm. Cells were maintained at 37°C during imaging using a Tokai Hit INUBG2E-ZILGS stage-top incubator. The microscope was additionally equipped with a p3E-300 illuminator (CoolLED), an MS-2000-XY motorized stage (ASI), and an LB10-3 filter wheel (ASI). pHluorin fluorescence was detected using a GFP filter set (Chroma, #49002) and excitation with a 491-nm, 100-mW Cobolt Calypso laser (Cobolt). pHmScarlet fluorescence was detected using a mCherry filter set (Chroma, #49008) and excitation with a 100-mW Cobolt Jive laser (Cobolt). Microscope acquisition was controlled using MetaMorph software version 7.10.4 (Molecular Devices).

MVB–plasma membrane fusion events were visualized using CD63-, CD9-, TSPAN2-, or TSPAN3-pHluorin fusion proteins. The fluorescence of pHluorin is quenched within the acidic lumen of intracellular compartments and rapidly increases upon fusion with the plasma membrane and exposure to the neutral extracellular environment. Cells were imaged at 300 ms/frame for 3 minutes in fresh DMEM. Fusion events were identified as transient, spatially confined increases in pHluorin fluorescence at the plasma membrane and quantified using the ExoU plugin in FIJI. Fusion-event frequency was expressed as events per cell.

For dual-color TIRF microscopy, cells co-expressing TSPAN3-pHluorin and CD63-pHmScarlet were imaged using the corresponding green and red fluorescence channels. Fusion events occurring within and 1 second between the two channels were classified as double-positive, whereas events detected exclusively in one channel were classified as TSPAN3-or CD63-positive ExoU plugin in FIJI.

### 3D vesicle colocalization analysis

Three-dimensional colocalization of CD63-and TSPAN3-positive intracellular vesicles was analyzed using arivis Vision4D (ZEISS arivis). Z-stacks containing the complete cellular volume were analyzed in three dimensions without plane-wise segmentation.

CD63-and TSPAN3-positive vesicles were independently segmented using the Blob Finder algorithm. Following segmentation, detected objects were filtered based on their three-dimensional volume to exclude structures outside the predefined vesicle-size range. The resulting CD63-and TSPAN3-positive vesicle populations were subsequently compared using the arivis Compartmentalization analysis. Vesicles were classified as colocalized when the segmented objects showed at least 50% three-dimensional overlap.

The numbers of CD63-positive, TSPAN3-positive, and CD63/TSPAN3 double-positive vesicles were determined for each analyzed cell. The relative abundance of each vesicle population was calculated as a percentage of the total number of segmented CD63-and/or TSPAN3-positive vesicles.

### HALO surface labeling and internalization microscopy assay

Plasma membrane exposure and subsequent internalization of endogenous CD63-HALO and TSPAN3-HALO were assessed using sequential labeling with cell-impermeant and cell-permeant HALO ligands.

For pulse–chase experiments, cells were incubated with cell-impermeable HALO ligand (iJF635, Janelia Research Campus) at 1 µM for 1 or 60 minutes. Cells were subsequently washed extensively with DMEM to remove unbound ligand. The remaining HALO-tagged protein pool was labeled with a spectrally distinct cell-permeable HALO ligand (JFX554, Janelia Research Campus) at 244.4 nM for 1 min. Cells were subsequently washed extensively with DMEM to remove unbound ligand and incubated in ligand-free medium for 90 min at 37°C to allow internalization of the surface-labeled protein pool. Following the chase period, cells were fixed with 4% paraformaldehyde containing 4% sucrose.

Images were acquired using an inverted Zeiss LSM 900 confocal laser-scanning microscope (Carl Zeiss Microscopy) equipped with a Plan-Apochromat 63x/1.4 WD=0.19mm, Oil Immersion. Fluorescence channels were acquired sequentially using the appropriate excitation wavelengths and detector settings. Acquisition settings were kept constant between CD63-HALO and TSPAN3-HALO samples within individual experiments. Intracellular cell-impermeant HALO fluorescence was quantified using FIJI. The 1-min pulse followed by a 90-min chase was used as the primary readout of plasma membrane exposure followed by endocytic retrieval. The 60-min continuous-labeling condition was used as a higher-sensitivity measurement of cell-impermeant HALO labeling.

### HALO surface labeling and flow cytometry

Cell-surface exposure of endogenous CD63-HALO and TSPAN3-HALO was quantified using a cell-impermeant HALO ligand. Cells were trypsinized, resuspended in DMEM containing 1 µM cell-impermeable HALO ligand (iJF635, Janelia Research Campus), and incubated for 1 min. Cells were subsequently washed with DMEM followed by FACS buffer (PBS supplemented with 2% FBS) and resuspended in FACS buffer for flow cytometric analysis. Fluorescence was measured using a BD FACSymphony A1 Cell Analyzer (BD Biosciences). Flow cytometry data were analyzed using FlowJo software (version 10, BD Biosciences).

### Immunofluorescence

HeLa cells were fixed with 4% PFA containing 4% sucrose for 10 min at room temperature, washed with PBS, and permeabilized with 0.5% Triton X-100 in PBS for 30 min. Cells were subsequently blocked with 2% bovine serum albumin (BSA) in PBS for at least 30 minutes and incubated overnight at 4°C with primary antibodies against EEA1 (BD Biosciences, #610457, 1:300), LAMTOR4 (Cell Signalling, #12284, 1:300), LAMP1 (BD Biosciences, #555798, 1:300), LC3 (MBL international corporation, #PM036, 1:300), or GABARAPL2 (Proteintech, 18724-1-AP, 1:300) at the indicated dilutions. Following washing with PBS, cells were incubated with the appropriate fluorophore-conjugated secondary antibodies (ThermoFisher, anti-mouse-AF488, #A11001, anti-rabbit-AF488, #A11008, 1:500) for 2 h at room temperature. For visualization of endogenous HALO-tagged CD63 and TSPAN3, live cells were labeled with JFX650 HALO ligand (Janelia Research Campus) at 1 µM for 1 min at room temperature prior to fixation. Samples were mounted in Vectashield containing DAPI and imaged using an inverted Zeiss LSM 900 confocal laser-scanning microscope (Carl Zeiss Microscopy).

### Image analysis and colocalization

Fluorescence images were analyzed using FIJI/ImageJ. Image acquisition and analysis settings were kept identical between experimental conditions within individual experiments. Individual cells were defined using Cellpose-based segmentation, and subsequent analyses were restricted to the corresponding cellular ROIs. Background fluorescence was subtracted using the rolling-ball algorithm in FIJI.

For analysis of the subcellular localization of endogenous CD63-HALO and TSPAN3-HALO, the HALO channel and the corresponding EEA1, LAMTOR4, LAMP1, or LC3 channel were manually thresholded. Colocalization was quantified using Manders’ colocalization coefficient (M1/M2) after thresholding, representing the fraction of signal overlap between the HALO signal and the respective marker channel. The same thresholding and object-detection parameters were applied to all conditions within each experiment.

For analysis of LC3-positive compartments in cells expressing CD63 WT, TSPAN3 WT, or the indicated TSPAN3 mutants, an LC3 intensity threshold was determined from control-transfected cells. For each experiment, the median LC3 fluorescence intensity was determined for each control cell, and the threshold was defined as the mean of these median values plus four standard deviations. Colocalization between tetraspanin-positive structures and LC3-positive puncta was quantified in both directions using Manders’ colocalization coefficients (M1 and M2), where M1 represents the fraction of TSPAN-positive signal overlapping LC3-positive signal and M2 represents the fraction of LC3-positive signal overlapping TSPAN-positive signal. To quantify changes in the LC3-positive compartment, LC3 fluorescence intensity was measured within thresholded LC3-positive pixels for each cell. Integrated LC3 intensity within the thresholded area was calculated and expressed relative to the corresponding control condition.

For all quantitative imaging experiments, individual cells were analyzed from at least 3 independent biological experiments. Image analysis was performed using identical analysis parameters across conditions within each experiment.

### GFP affinity purification

GFP affinity purification was performed to identify proteins associated with CD63-and TSPAN3-pHluorin and to assess changes in the TSPAN3-associated proteome upon mutation of selected cytosolic residues. HeLa cells expressing CD63-pHluorin, TSPAN3-pHluorin, GFP control, or the indicated TSPAN3-pHluorin mutants were cultured in a 6-well plate and harvested 24 hours after transfection. Cells were washed with ice-cold PBS and lysed in 1x RIPA buffer (50mM Tris-HCL pH 7.4; 150mM NaCl; 1% Triton X-100; 0.5% Sodium Deoxycholate; 0.1% SDS; 1mM EDTA; 10mM NaF) supplemented with 1x Protease Inhibitor Cocktail (Roche; #11836170001) for 30 min on ice.

Cell lysates were cleared by centrifugation at 16100×*g* for 15 min at 4°C. Protein concentrations were determined micro BCA™ protein assay kit (Thermo Scientific, #23235) and equal amounts of total protein were used for each affinity purification. Cleared lysates were incubated with GFP-Trap agarose (Proteintech, #gta) overnight at 4°C with end-over-end rotation. Beads were subsequently collected and washed 5 times with ice-cold PBS to remove non-specifically associated proteins.

After the last wash, all supernatant was removed using a 25g 0.5 x 16 mm needle. Elution buffer (2M Urea, 100 mM Tris pH 7.5, 12 mM DTT) was and beads were incubated at RT at maximum speed on a shaker for 20 minutes. Next, iodoacetamide was added to a final concentration of 50 mM followed by incubation on a shaker in the dark for 10 minutes. 0.1 µg trypsin was added and samples were incubated for 2 hours on a shaker at maximum speed. Samples were centrifuged at 1500 x *g* for 2 minutes at RT and the supernatant containing the eluted peptides was transferred to a new PCR tube. To ensure all polypeptides were collected from the beads, elution buffer was added once more and beads were incubated for 5 minutes at RT. The tubes were centrifuged at 1500 x *g* for 2 minutes, and the supernatant was combined with the eluate retrieved in the previous step. 0.033 µg trypsin was added to the solution and tubes were incubated on a shaker at RT overnight. After overnight incubation, polypeptides were purified using StageTips^36^. Two layers of C18 (cat. 66883-U, Supelco) were inserted into a P200 pipette tip to make the StageTip, which were washed once with ULC/MS grade methanol (Biosolve), once with buffer B (0.1% (v/v) ULC/MS grade formic acid (Biosolve) and 80% (v/v) acetonitrile (VWR) in ultrapure water), and twice with buffer A (0.1% (v/v) ULC/MS grade formic acid (Biosolve) in ultrapure water) using centrifugation at 1500 x *g* for 4 minutes at RT. Polypeptides were acidified by addition of a final concentration of 1% trifluoroacetic acid (TFA, Biosolve) and loaded on the washed StageTips. After centrifugation at room temperature at 600 x *g* for 10 minutes, StageTips were washed with buffer A using centrifugation at 600 x *g* at room temperature for 2 minutes. For mass spectrometry analysis, peptides were eluted from the StageTips and analyzed on an Orbitrap Astral mass spectrometer (ThermoFisher Scientific). Data was collected in data-independent acquisition (DIA) mode using 24-minute gradients. Raw mass spectrometry spectra were analyzed per cell type using DIA-NN with match between runs enabled^37^. Proteins were identified using a search against the human UniProtKB human proteome (downloaded 5 June 2024), taking along contaminants. A maximum number of two missed cleavages and five variable modifications were allowed. Methionine oxidation (15.9949 Da) and N-terminal acetylation (42.0106 Da) were added as dynamic modifications. MS2 was set to 10 ppm and MS1 to 4 ppm and the optimal scan window was determined individually for each experiment. The resulting protein identification file was filtered for contaminants, reproducibly identified proteins, and missing values were imputed from the normal distribution^38^. Groups were compared using a two sample t-test (S0 = 2, False Discovery Rate < 0.05).

For identification of the TSPAN3-associated proteome, proteins enriched with TSPAN3-pHluorin were compared with the GFP control and with CD63-pHluorin. To determine the contribution of individual TSPAN3 residues to its associated protein network, GFP affinity purification was additionally performed using TSPAN3(T75A), TSPAN3(Y252A), TSPAN3(LL246AA), and TSPAN3(Y244A)-pHluorin, and their associated proteomes were compared with TSPAN3 WT.

### ultraID proximity biotinylation and streptavidin enrichment

The proximity proteomics data analyzed in this study were generated previously and correspond to the same dataset reported by Zala et al^17^ and were reanalyzed in the present study to compare the molecular environments of CD63 and TSPAN3. Briefly, HeLa cells expressing Myc-ultraID-CD63, Myc-ultraID-TSPAN3 were cultured in 15-cm dishes and incubated with 50 µM biotin to induce proximity-dependent labeling of proteins surrounding the respective tetraspanin. Wild-type HeLa cells subjected to the same biotin-labeling conditions were included as controls.

For visualization of proximity-labeling activity by fluorescence microscopy, cells were incubated with biotin for 30 min. Cells were subsequently washed with PBS, fixed with 4% PFA, permeabilized with 0.5% Triton X-100 in PBS, blocked with 2% BSA in PBS and stained with an mouse-anti-Myc antibody (Santa Cruz Biotechnologies, sc-40, 1:500) and anti-mouse-AF488 secondary antibody (ThermoFisher #A11001, 1:500) to visualize the ultraID fusion proteins. Biotinylated proteins were detected using AlexaFluore-555 conjugated streptavidin (ThermoScientific, S21381, 1:1000). Samples cultured in the absence of exogenous biotin were included as controls.

For proximity proteomics, cells expressing Myc-ultraID-CD63 or Myc-ultraID-TSPAN3 were incubated with biotin for 24 h. Following labeling, cells were washed extensively with ice-cold PBS to remove extracellular biotin and harvested. Wild-type HeLa cells subjected to the same biotin-labeling conditions were included as controls. Cell pellets were lysed in RIPA buffer and protein concentrations were determined by micro BCA™ protein assay kit (Thermo Scientific, #23235).

Biotinylated proteins in this previously generated dataset were enriched using Neutravidin affinity purification as described by Zala et al^17^. Briefly, equal amounts of protein were incubated with 20 µL Pierce Neutravidin Agarose beads (Thermo Fisher Scientific) for 4 h at room temperature using an overhead rotator. Following incubation, beads were collected by centrifugation and subjected to sequential washing steps to remove non-specifically associated proteins. Enriched proteins were digested on-bead with trypsin (Promega, V5111; enzyme-to-protein ratio 1:25, 0.2 µg trypsin per sample) for 1 h at 47 °C. The resulting peptides were analyzed by liquid chromatography–tandem mass spectrometry (LC–MS/MS).

### Biotinylated EV

The biotinylated extracellular vesicle (EV) proteomics data analyzed in this study correspond to the dataset previously generated and reported by Zala et al^17^. and were reanalyzed here to compare proteins associated with CD63-and TSPAN3-positive EVs. HeLa cells expressing Myc-ultraID-CD63 or Myc-ultraID-TSPAN3 were incubated with 50 µM biotin for 24 h to induce proximity-dependent biotinylation. Wild-type HeLa cells subjected to the same biotin-labeling conditions were included as controls.

Following labeling, conditioned medium was collected and extracellular vesicles were isolated by size-exclusion chromatography (SEC) as previously described^17^. EV-containing fractions were pooled and concentrated prior to protein extraction. Biotinylated proteins present in the isolated EV preparations were subsequently enriched using NeutrAvidin affinity purification. Equal amounts of EV-derived protein were incubated with Pierce NeutrAvidin Agarose beads (Thermo Fisher Scientific), followed by sequential washing steps to remove non-specifically associated proteins.

Enriched proteins were digested on-bead with trypsin and the resulting peptides were analyzed by liquid chromatography–tandem mass spectrometry (LC–MS/MS) as previously described17. The resulting proteomic dataset was used in the present study for comparative analysis of proteins enriched in CD63-and TSPAN3-associated EVs.

### Proteomics analysis

Gene set enrichment analysis (GSEA) was performed using the GSEA Desktop application and gene sets from the Molecular Signatures Database (MSigDB)^18^. For the TSPAN3-pHluorin versus GFP-control affinity-purification proteomics dataset, identified proteins were ranked according to their signed t-statistic and analyzed using the GSEAPreranked module. Enrichment was assessed separately against the MSigDB C5 Gene Ontology Biological Process (GO:BP), C5 Gene Ontology Cellular Component (GO:CC), and C2 Reactome collections. Analyses were performed using 1,000 gene-set permutations, weighted enrichment scoring, and gene-set size limits of 15–500 genes. Enrichment was reported as normalized enrichment scores (NES) with corresponding nominal *P* values and false discovery rate (FDR) q-values.

### Electron microscopy

Cells in culture were initially fixed for 5 min with 2% paraformaldehyde (PFA) in 0.1 M phosphate buffer mixed 1:1 with culture medium, followed by fixation with fresh 4% PFA in the same buffer for 2 h at room temperature. After washing in PBS, residual aldehydes were quenched with 0.15% glycine in PBS for 10 min. Quenched cells were scraped, resuspended in 1% gelatin/PBS, and infiltrated with 12% gelatin at 37 °C for 10 min. Cell pellets were solidified on ice for 30 min, trimmed into small blocks, and cryoprotected overnight in 2.3 M sucrose at 4 °C before storage in liquid nitrogen. Sucrose-infused blocks were sectioned at 80 nm (−110 °C) using a diamond knife on a Leica ultracryomicrotome. Sections were picked up on Formvar/carbon-coated grids using a 1:1 mixture of 2.3 M sucrose and 1.8% methylcellulose. Grids were incubated at 37 °C for 30 min to remove gelatine, blocked with 0.5% fish skin gelatine and 0.1% BSA-c in PBS, and labelled with a primary antibody against GFP (Rockland rabbit anti-GFP, 1:200). Detection was performed using Protein A conjugated to 10-nm colloidal gold (CMC, UMC Utrecht). Labeled sections were post-fixed in 1% glutaraldehyde, washed in Milli-Q water, and contrasted on ice using 2% uranyl oxalate (pH 7.0, 5 min) followed by uranyl acetate/methylcellulose (pH 4.0, 10 min). Grids were dried using the loop-out method and imaged on a Tecnai 12 TEM operated at 80 kV^19,20^.

## Results

### TSPAN3 marks a distinct secretory MVB-population

Extracellular vesicles can originate from distinct cellular compartments, including the plasma membrane and the endosomal system. Because multivesicular bodies (MVBs) can either undergo lysosomal degradation or fuse with the plasma membrane to release intraluminal vesicles (ILVs), we first asked whether different tetraspanin-defined EV populations differ in their dependence on endolysosomal trafficking (Figure 1A). To quantify the release of CD63-, CD9-, TSPAN2-and TSPAN3-positive EVs, we used a NanoLuciferase (Nluc)-based EV secretion assay^16^. HA-Nluc was knocked-in at the N-terminus of the endogenous tetraspanins using CRISPR/Cas9-mediated genome editing (Supplementary Figure 1A–C). Size-exclusion chromatography of conditioned medium showed that extracellular Nluc activity co-eluted with EV-containing fractions, supporting its use as a reporter of EV-associated secretion (Supplementary Figure 1D). Inhibition of lysosomal acidification with Bafilomycin A1 (BafA1) significantly increased EV secretion of HA-Nluc-CD63 and HA-Nluc-TSPAN3, whereas secretion of HA-Nluc-CD9 and HA-Nluc-TSPAN2 was not significantly affected (Figure 1B, Supplementary Figure 1E, F). Notably, the BafA1-induced increase was greater for TSPAN3 than for CD63 (8.2-fold versus 3.5-fold, P = 0.0185). These findings indicate that TSPAN3-positive EV secretion is strongly coupled to endolysosomal trafficking and suggest that TSPAN3 preferentially labels an MVB-derived EV population.

**Figure 1:**
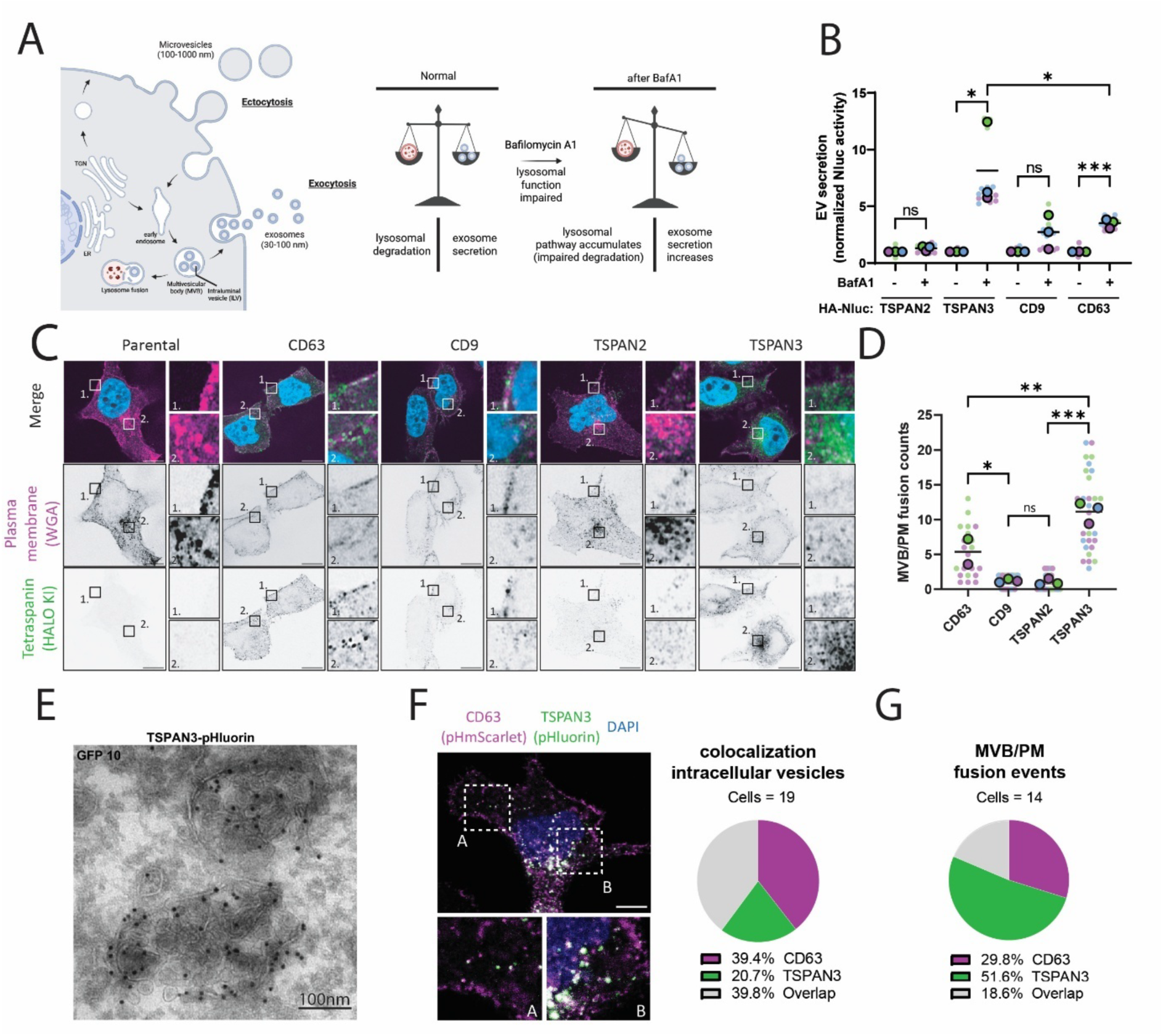
TSPAN3 marks a distinct population of secretory MVBs. (A) Schematic overview of extracellular vesicle (EV) biogenesis pathways, including plasma membrane-derived microvesicles, and multivesicular body (MVB)-derived exosomes. Bafilomycin A1 (BafA1) inhibits lysosomal acidification and degradation, thereby altering the balance between lysosomal degradation and secretion of MVB-derived EVs.(B) Nluc-based quantification of EV secretion from HA-Nluc-TSPAN2, HA-Nluc-TSPAN3, HA-Nluc-CDG, and HA-Nluc-CD63 knock-in cells in the absence or presence of BafA1. (C) Representative super-resolution images showing the cellular localization of endogenous HALO-TSPAN2, HALO-TSPAN3, HALO-CDG, and HALO-CD63. HALO-tagged proteins were labeled with CA-MaP555 and the plasma membrane was visualized using wheat germ agglutinin (WGA). (D) Quantification of MVB–plasma membrane fusion events detected by TIRF microscopy using TSPAN2-pHluorin, TSPAN3-pHluorin, CDG-pHluorin, and CD63-pHluorin reporters. (E) Representative immunogold electron micrograph of TSPAN3-pHluorin showing localization of pHluorin immunoreactivity within an MVB. Scalebar is 100nm. (F) Dual-color fluorescence microscopy of CD63-pHmScarlet and TSPAN3-pHluorin showing the relative proportions of CD63-positive, TSPAN3-positive, and double-positive intracellular vesicles. (G) Dual-color TIRF microscopy of CD63-pHmScarlet and TSPAN3-pHluorin showing the relative proportions of CD63-positive, TSPAN3-positive, and double-positive MVB–plasma membrane fusion events. Data are from n = 3 independent experiments. Each big point represents the mean of an individual experiment. Each small point represents an individual quantification. Statistical significance was determined using one-way anova; ns, not significant; *P < 0.01, *P* < 0.001, *P < 0.0001. Scale bars, 20 µm unless otherwise indicated.

We next asked whether TSPAN3 localizes to intracellular compartments consistent with an endosomal origin. To determine the subcellular localization of these EV populations, we introduced HALO into the endogenous loci of the first extracellular loops of CD63, CD9, TSPAN2 and TSPAN3 using CRISPR/Cas9-mediated genome editing. Super-resolution microscopy following labeling with the fluorescent HALO ligand CA-MaP555 and the plasma membrane marker WGA revealed distinct localization patterns (Figure 1C). CD9 and TSPAN2 localized primarily to the plasma membrane, whereas TSPAN3 was predominantly associated with intracellular vesicular distribution. In contrast, CD63 was detected both at the plasma membrane and on intracellular vesicles.

We then asked whether these intracellular TSPAN3-positive compartments represent secretory MVBs. To directly visualize membrane fusion, we performed TIRF microscopy using pH-sensitive tetraspanin-pHluorin reporters (Figure 1D)^15^. Frequent fusion events were detected for TSPAN3-pHluorin and CD63-pHluorin, whereas substantially fewer events were observed for CD9-pHluorin and TSPAN2-pHluorin. TSPAN3-pHluorin exhibited a higher frequency of fusion events than CD63-pHluorin. Consistent with an endosomal origin, immunogold electron microscopy localized TSPAN3-pHluorin to MVBs and their intraluminal vesicles (Figure 1E, Supplementary Figure 1G). Together, these observations identify TSPAN3 as a component of ILV-containing MVBs that undergo plasma membrane fusion.

Finally, we asked whether TSPAN3 marks the same secretory MVB population as the canonical EV marker CD63. Analysis of intracellular vesicles in cells co-expressing CD63-pHmScarlet and TSPAN3-pHluorin showed that 39.8% were double-positive, whereas 38.4% and 20.7% were exclusively positive for CD63 and TSPAN3, respectively (Figure 1F). We subsequently used dual-color TIRF microscopy to determine the composition of MVBs undergoing plasma membrane fusion. In contrast to the intracellular pool, only 18.6% of fusion events contained both CD63 and TSPAN3, whereas 29.8% were CD63-only and 51.6% were TSPAN3-only (Figure 1G, Supplementary Figure 1H,I). Thus, TSPAN3-only compartments were strongly enriched within the secretory MVB population, while CD63/TSPAN3 double-positive compartments were comparatively underrepresented. Together, these findings establish TSPAN3 as a marker of a secretory MVB population that is largely distinct from canonical CD63-positive compartments, providing evidence that molecularly distinct MVB populations contribute differentially to EV secretion.

### TSPAN3 reaches secretory MVBs through a trafficking route distinct from CD63

Having established that TSPAN3 and CD63 mark partially distinct populations of secretory MVBs, we next asked whether they reach these compartments through distinct intracellular trafficking routes. CD63 is known to traffic to late endosomal compartments in part through delivery to the plasma membrane followed by endocytic retrieval^6^. We therefore asked whether TSPAN3 similarly depends on endocytosis to access secretory MVBs.

We first asked whether CD63-and TSPAN3-positive EV secretion respond similarly to perturbation of endocytic trafficking. Inhibition of dynamin-dependent endocytosis with dynasore markedly reduced EV-associated CD63 and also decreased TSPAN3 secretion, although the reduction was less pronounced for TSPAN3 (Fig. 2A). Cellular levels of both proteins were maintained or increased following dynasore treatment, indicating that the reduction in EV-associated signal was not explained by decreased protein abundance. In contrast, treatment with the clathrin-terminal-domain inhibitor Pitstop 2 did not significantly alter EV secretion or cellular levels of either CD63 or TSPAN3 (Fig. 2B). Together, these pharmacological experiments indicated that endocytic trafficking contributes to secretion of both tetraspanins, while suggesting that TSPAN3 may be less dependent on this route than CD63. This prompted us to investigate the molecular basis of these differences.

We next asked whether CD63 and TSPAN3 rely on the same intracellular sorting determinants. Similar to CD63, the C-terminal tail of TSPAN3 contains a YXXΦ-motif predicted to interact with adaptor protein complexes, as well as an adjacent dileucine-based sorting motif. (Fig. 2C). The TSPAN3 YXXΦ-containing region has recently been shown to interact with AP2, whereas its dileucine motif regulates intracellular localization in human B cells^21^. We therefore initially hypothesized that CD63 and TSPAN3 might reach MVBs through a similar adaptor-dependent trafficking mechanism. To address this, we generated mutants dirsupting the CD63 YXXΦ motif and TSPAN3-YXXΦ and dileucine motifs. Mutation of the CD63 YXXΦ motif resulted in pronounced redistribution toward the plasma membrane and reduced intracellular localization (Fig. 2D). Consistent with this redistribution, the frequency of CD63-positive MVB–plasma membrane fusion events was reduced by 3.28-fold (to ∼30% of wild-type levels) compared with wild-type CD63 (Fig. 2E).

Unexpectedly, disruption of the corresponding YXXΦ motif in TSPAN3 had a substantially smaller effect. The TSPAN3 YXXΦ mutant retained prominent intracellular localization (Fig. 2D), and MVB–plasma membrane fusion events were reduced to approximately 56% of wild-type levels, a difference that did not reach statistical significance (Fig. 2E). By contrast, mutation of the adjacent TSPAN3 dileucine motif strongly altered its localization and reduced fusion events to approximately 11% of wild-type levels (Fig. 2D,E). Together, these findings indicate that CD63 and TSPAN3 use distinct sorting determinants to access secretory MVBs. Whereas CD63 primarily depends on its YXXΦ motif, selective disruption of the TSPAN3 YXXΦ tyrosine had only a modest effect, while mutation of the overlapping LL residues strongly impaired trafficking to secretory MVBs. This suggests that determinants associated with the LL-containing region, rather than canonical YXXΦ-dependent sorting, make the dominant contribution to TSPAN3 trafficking.

The distinct sorting requirements suggested that CD63 and TSPAN3 may also differ in their intracellular trafficking routes. We therefore asked whether TSPAN3, like CD63, traffics through the plasma membrane before reaching intracellular compartments. To address this, we compared plasma membrane exposure and endocytic retrieval of endogenous CD63 and TSPAN3 using HALO tagging using cell-impermeable HALO ligands. Flow-cytometric analysis revealed substantially lower surface labeling of TSPAN3-HALO than CD63-HALO, indicating that a smaller fraction of the TSPAN3 pool is exposed at the plasma membrane at steady state (Fig. 2F). To determine whether this difference was accompanied by reduced endocytic uptake, surface-exposed CD63-HALO and TSPAN3-HALO were pulse-labeled with cell-impermeable HALO ligand and followed during internalization (Fig. 2G). CD63 showed pronounced accumulation of internalized surface-derived signal over 60 min, whereas internalization of TSPAN3 was substantially lower (Fig. 2G,H). Although prolonged labeling may also permit uptake of extracellular ligand into intracellular compartments, quantification nevertheless revealed a 2.18-fold lower intracellular signal for TSPAN3 than for CD63 after 60 min (P = 0.045). Together, these data indicate that plasma membrane exposure followed by endocytic retrieval is a prominent trafficking route for CD63, but contributes substantially less to the intracellular TSPAN3 pool.

Together, these findings demonstrate that TSPAN3 reaches secretory MVBs through a trafficking route that is mechanistically distinct from that of CD63 (Fig. 2I). CD63 undergoes extensive plasma membrane exposure followed by YXXΦ-dependent endocytic retrieval, whereas only a limited fraction of TSPAN3 is detectably exposed at the plasma membrane. Instead, TSPAN3 depends predominantly on its dileucine motif for delivery to fusion-competent secretory MVBs, indicating that it primarily reaches these compartments through an intracellular trafficking route while retaining a smaller contribution from plasma membrane endocytosis.

**Figure 2:**
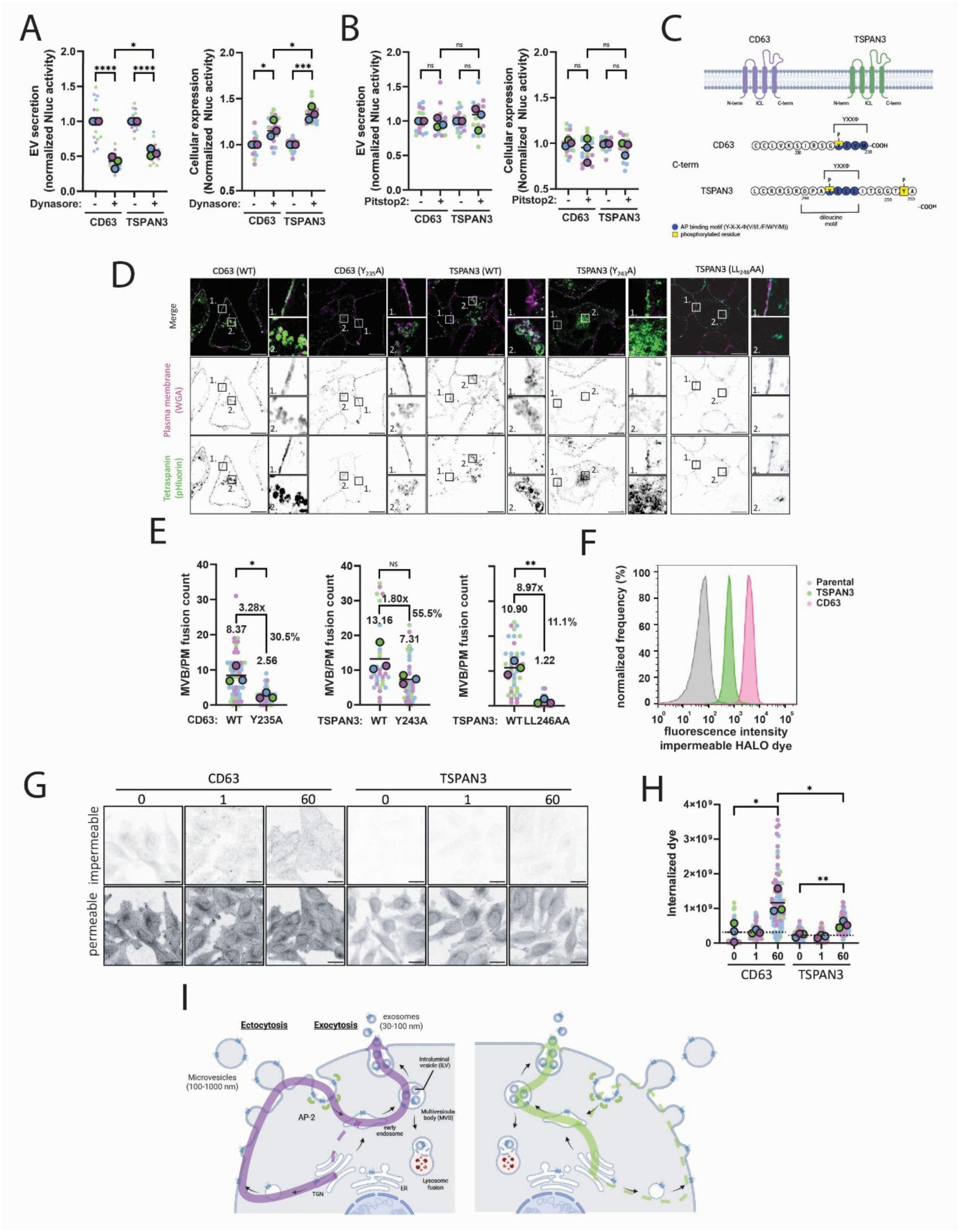
TSPAN3 reaches secretory MVBs through a trafficking route distinct from CD63. (A, B) Nluc-based quantification of EV secretion from HA-Nluc-CD63 and HA-Nluc-TSPAN3 cells following treatment with the endocytic trafficking inhibitors dynasore (A) or Pitstop 2 (B). (C) Schematic overview of CD63 and TSPAN3 C-terminal YXXΦ-and dileucine-motifs. (D) Representative super-resolution fluorescence images of wild-type and YXXΦ/dileucine trafficking mutants of CD63 and TSPAN3. The plasma membrane was visualized using wheat germ agglutinin (WGA). Insets show magnified regions indicated in the merged images. (E) Quantification of MVB/plasma membrane fusion events for wild-type and YXXΦ/dileucine mutants of CD63 and TSPAN3 using pHluorin-based TIRF microscopy. Values above the indicated conditions denote mean MVB/PM fusion count. (F) Histogram showing membrane expression of parental (gray), TSPAN3-HALO (green) and CD63-HALO (magenta) (G) Representative confocal images from a HALO pulse-chase assay in which surface-exposed CD63-HALO and TSPAN3-HALO were labeled with a cell-impermeable HALO ligand, followed by labeling of the remaining protein pool with a cell-permeable HALO ligand. (H) Quantification of internalized cell-impermeable HALO signal following the pulse-chase assay shown in (G). (I) Working model illustrating predominant trafficking routes of CD63 (magenta/left) and TSPAN3 (green/right) to secretory MVBs. CD63 undergoes substantial plasma membrane delivery and endocytic retrieval before reaching MVBs, whereas TSPAN3 predominantly reaches MVBs through an intracellular trafficking route with limited plasma membrane transit. Data are from n = 3 independent experiments. Each big point represents the mean of an individual experiment. Each small point represents an individual quantification. Statistical significance was determined using one-way ANOVA; ns, not significant; *P < 0.01, *P* < 0.001, *P < 0.0001. Scale bars, 20 µm unless otherwise indicated.

### Proteomics identifies a distinct molecular machinery associated with TSPAN3

Having established that TSPAN3 reaches secretory MVBs through a trafficking route distinct from CD63, we next asked whether this difference is reflected in the molecular machinery associated with the two tetraspanins. We performed ultraID-based proximity biotinylation followed by quantitative mass spectrometry (Figure 3A)^17^. Myc-ultraID was fused to either CD63 or TSPAN3, enabling biotinylation of proteins within the local environment of each tetraspanin. To confirm proximity-labeling activity, cells expressing Myc-ultraID-CD63 or Myc-ultraID-TSPAN3 were incubated with biotin and analyzed by fluorescence microscopy. Addition of biotin resulted in a pronounced increase in streptavidin staining surrounding the ultraID fusion proteins, confirming efficient proximity biotinylation (Figure 3B; Supplementary Figure 2A).

We first asked which cellular compartments and molecular processes are represented within the TSPAN3 molecular environment. Following 24 h of biotin labeling, biotinylated proteins were enriched by streptavidin affinity purification and analyzed by mass spectrometry. Both Myc-ultraID-CD63-and Myc-ultraID-TSPAN3-expressing cells showed enrichment of biotinylated proteins compared with the ultraID control condition (Supplementary Figure 2B,C). To obtain an unbiased overview of the TSPAN3 molecular environment, we performed gene set enrichment analysis using proteins ranked by their enrichment in TSPAN3-ultraID relative to the control. Gene Ontology Cellular Component analysis revealed strong enrichment of membrane-trafficking compartments and complexes, including the SNARE complex, multivesicular body membrane, ESCRT complex, endosomal and late-endosomal membranes, autophagosome membrane, and exocytic vesicles (Figure 3C). Thus, the TSPAN3 molecular neighborhood was strongly associated with endosomal, MVB, autophagic, and membrane-fusion machinery.

We next asked which proteins were preferentially enriched in the TSPAN3 molecular environment relative to CD63. Direct comparison of the CD63 and TSPAN3 proximity proteomes identified 38 proteins significantly enriched in the proximity of TSPAN3 relative to CD63 (Figure 3D; Supplementary Figure 2D). These included proteins involved in intracellular membrane trafficking and the endolysosomal system, such as the adaptor-complex component AP1M1, the ESCRT-I component VPS37A, the phosphoinositide phosphatase FIG4, the vesicular trafficking GTPase RAB3D, and SDCBP/syntenin-1, a well-established regulator of endosomal trafficking and exosome biogenesis. Several TSPAN3-enriched proteins were also linked to lysosomal signaling and autophagy regulation, including TFEB, FLCN, and LAMTOR3. Together with the gene set enrichment analysis, these data indicate that although TSPAN3 and CD63 share much of their molecular environment, TSPAN3 is preferentially associated with a subset of established endolysosomal and EV biogenesis regulators, as well as proteins linked to autophagy.

To determine whether these associations could be confirmed using an orthogonal biochemical approach, we performed GFP affinity purification followed by mass spectrometry using lysates from TSPAN3-pHluorin-expressing cells (Figure 3E). TSPAN3-pHluorin was enriched using an anti-GFP affinity matrix, and co-purified proteins were compared with a GFP control. Quantitative proteomic analysis identified a distinct set of proteins significantly enriched in the TSPAN3-pHluorin pulldown (Supplementary Figure 2F).

We next asked whether the two complementary proteomic approaches converged on a common TSPAN3-associated molecular network. We compared protein enrichment in the TSPAN3 affinity-purification and proximity-labeling datasets (Figure 3F). Despite the different experimental principles of the two approaches, several proteins associated with endosomal, lysosomal, and vesicular trafficking were enriched in both datasets. Among these was SDCBP/syntenin-1, a regulator of endosomal sorting and intraluminal vesicle formation linked to exosome biogenesis. Additional shared hits included the lysosomal membrane proteins TMEM9, SLC38A9, MFSD8, and SIDT2, as well as the endolysosomal ubiquitin ligase RNF167 and the membrane-trafficking protein SCAMP2. The adaptor-complex component AP2M1 was also enriched in both datasets. Notably, the ATG8-family protein GABARAPL2 was reproducibly enriched by both approaches, further suggesting that TSPAN3-associated membranes intersect with autophagy-related machinery.

Inspection of the TSPAN3 proximity proteome further revealed a prominent group of proteins involved in late-endosomal and autophagic membrane fusion. These included RAB7A, STX17, SNAP29, and YKT6, together with the late-endosomal SNAREs STX7, VAMP7, and VTI1B (Figure 3F). RAB7A is a central regulator of late-endosome maturation, whereas STX17 and SNAP29 participate in SNARE-dependent autophagosome–lysosome fusion. YKT6 has similarly been implicated in autophagic membrane fusion and was among the most strongly enriched proteins in the TSPAN3 proximity proteome. The coordinated enrichment of these proteins was therefore consistent with the strong SNARE-and autophagosome-associated signatures identified by GSEA.

A second prominent module comprised proteins involved in ILV biogenesis and EV secretion. Indeed, the TSPAN3 proximity proteome also contained multiple components of the ESCRT machinery, including HGS, STAM2, VPS25, CHMP4B, CHMP2B, IST1, and VPS4A/B, together with the MVB docking regulator RAB27A (Figure 3F). Their enrichment was consistent with the MVB membrane, ESCRT, endosomal membrane, and exocytic vesicle signatures identified by GSEA.

Together, these complementary proteomic approaches define the molecular environment of TSPAN3 and reveal an unexpectedly tight association with endosomal trafficking, MVB biogenesis, membrane fusion, EV secretion, and autophagy-related membrane remodeling. The recurrent identification of autophagy-associated proteins and compartments, including GABARAPL2, autophagosome-associated gene sets, and multiple components of autophagic membrane-fusion machinery, prompted us to investigate whether TSPAN3-containing compartments are functionally connected to the autophagy pathway.

**Figure 2.**
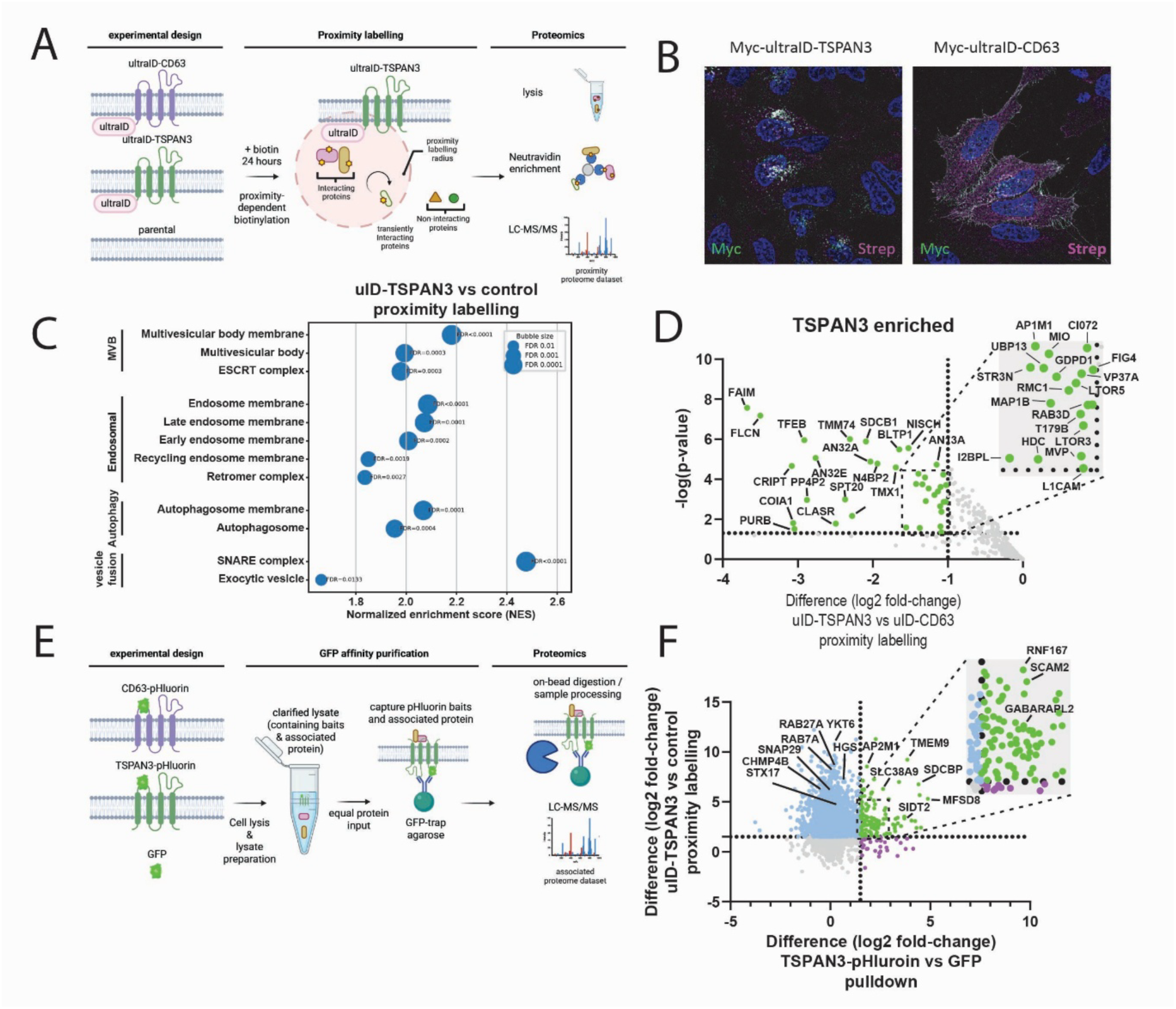
Orthogonal proteomic approaches identify a distinct TSPAN3-associated protein network. (A) Schematic overview of the ultraID-based proximity biotinylation approach used to identify proteins in the molecular proximity of TSPAN3 and CD63. Following addition of biotin, ultraID fused to the indicated tetraspanin biotinylates proximal proteins, which were subsequently enriched and identified by mass spectrometry. (B) Representative fluorescence images of cells expressing Myc-ultraID-TSPAN3 or Myc-ultraID-CD63 following 30 min of biotin labeling. Myc-ultraID fusion proteins were detected by anti-Myc staining (green), biotinylated proteins were visualized using fluorescent streptavidin (magenta), and nuclei are shown in blue. Scale bars, 20 µm. (C) Gene set enrichment analysis (GSEA) of the TSPAN3-ultraID proximity proteome relative to the ultraID control, showing selected enriched Gene Ontology Cellular Component terms. Normalized enrichment scores (NES) are shown; dot size indicates the false discovery rate (FDR). (D) Comparative analysis of the ultraID proximity proteomes of TSPAN3 and CD63. Proteins preferentially enriched in the proximity of TSPAN3 relative to CD63 are highlighted, with selected TSPAN3-enriched proteins indicated. (E) Schematic overview of the complementary GFP affinity purification–mass spectrometry approach using TSPAN3-pHluorin as bait. GFP-associated protein complexes were isolated by affinity purification and analyzed by mass spectrometry. (F) Comparison of protein enrichment obtained using the two independent proteomic approaches. Enrichment in the TSPAN3-ultraID proximity proteome relative to the ultraID control is plotted against enrichment in the TSPAN3-pHluorin pulldown relative to the GFP control. Proteins enriched in both approaches are highlighted, with selected proteins indicated. Proteomic analyses were performed from n = 3 independent biological replicates. Dashed lines indicate the fold-enrichment and adjusted P-value thresholds used to define enriched proteins in the respective comparisons.

### TSPAN3 engages autophagy-related machinery during trafficking to secretory MVBs

The TSPAN3-associated proteome identified in Figure 3 contained multiple proteins linked to endolysosomal trafficking and autophagy-related processes, suggesting that TSPAN3-positive compartments may intersect with autophagic membrane trafficking. We therefore first asked whether TSPAN3 localizes to autophagy-associated membrane compartments in cells. Endogenous CD63-HALO and TSPAN3-HALO were co-stained with EEA1, LAMTOR4, LAMP1, or LC3 to visualize early endosomes, late endosomal/lysosomal compartments, lysosomes, and autophagy-related membranes, respectively (Figure 4A). Quantification of colocalization revealed distinct compartmental distributions of CD63 and TSPAN3 (Figure 4B). CD63 showed greater association with EEA1, whereas TSPAN3 showed greater association with LamTOR4. Notably, TSPAN3 displayed a pronounced association with LC3-positive compartments compared with CD63, indicating that TSPAN3-positive membranes preferentially intersect with LC3-associated compartments.

We next asked whether this cellular association with LC3-positive compartments was also reflected in the TSPAN3-associated molecular environment. We therefore re-examined the TSPAN3-pHluorin GFP affinity-purification dataset for proteins involved in autophagy and autophagy–endosomal crosstalk (Figure 4C). TSPAN3 associated with proteins representing several stages of autophagy-related membrane trafficking. Notably, the ATG8-family protein GABARAPL2 was enriched in the TSPAN3 pulldown, together with proteins involved in selective autophagy, including NBR1, BNIP3 and CCPG1, and several ER-phagy-associated proteins, including RETREG1, RTN3 and TEX264. Particularly strong enrichment was observed for TMEM59, TMEM9 and RNF13, proteins positioned at the interface between endosomal and autophagy-related trafficking. In contrast, most components of the core autophagy initiation and conjugation machinery showed little or no enrichment. Together, these findings indicate that TSPAN3 is not broadly associated with the canonical autophagy machinery but instead preferentially associates with a selective network of ATG8-family, selective-autophagy, and endolysosomal trafficking proteins.

The prominent enrichment of GABARAPL2, together with the increased localization of TSPAN3 to LC3-positive compartments, raised the possibility that TSPAN3 contains sequence determinants that mediate its association with ATG8-family proteins. We therefore next asked whether specific cytosolic motifs in TSPAN3 contribute to this interaction. Inspection of the cytosolic regions of TSPAN3 identified the Y243 and LL246 as candidate LIR-associated sequence elements. In addition, two regions containing putative phosphorylation domains (T75 and Y252) were selected as candidate regulatory residues (Figure 4D). To examine their contribution to TSPAN3 trafficking and its association with autophagy-related machinery, we generated TSPAN3(T75A) and TSPAN3(Y252A) mutants. The previously characterized Y243A and LL246AA trafficking mutants were included as additional controls. Super-resolution imaging showed that both TSPAN3(T75A) and TSPAN3(Y252A) retained a predominantly intracellular vesicular distribution, indicating that mutation of these residues did not grossly disrupt TSPAN3 localization (Figure 4E). This contrasted with the pronounced plasma membrane redistribution observed for the LL246AA trafficking mutant described in Figure 2.

We next asked whether these mutations altered the molecular environment of TSPAN3. GFP affinity purification followed by quantitative mass spectrometry was performed for TSPAN3(T75A), TSPAN3(Y252A), and the previously characterized Y243A and LL246AA mutants, and changes in selected TSPAN3-associated proteins were compared with wild-type TSPAN3 (Figure 4F). The T75A mutant showed the most pronounced loss of autophagy-and endosomal-associated proteins, including a reduction in GABARAPL2 association, whereas the Y252A mutant showed a partially overlapping but distinct loss of associated proteins. In contrast, the Y243A mutant had relatively little effect on this interaction profile, while the LL246AA mutant produced a different pattern of changes. Together, these findings indicate that T75, and to a lesser extent Y252, contribute to the association of TSPAN3 with a selective autophagy-and endolysosomal protein network.

We therefore next asked whether disruption of these residues also altered the association of TSPAN3 with LC3-positive compartments. Cells expressing CD63 WT, TSPAN3 WT, TSPAN3(LL246AA), TSPAN3(T75A), TSPAN3(Y243A), or TSPAN3(Y252A) were stained for endogenous LC3 (Figure 4G). TSPAN3 WT showed a pronounced association with LC3-positive puncta compared with CD63, whereas this association was reduced for TSPAN3(T75A) and TSPAN3(Y252A), but remained largely unaffected by the LL246AA and Y243A mutations (Figure 4H,I). Thus, T75 and Y252 selectively contribute to the association of TSPAN3 with LC3-positive compartments. We next asked whether TSPAN3 not only associates with LC3-positive structures, but also alters the abundance or organization of these compartments. Expression of TSPAN3 WT increased LC3 signal compared with control or CD63-expressing cells (Figure 4J). This increase was significantly reduced by the T75A mutation and showed a decreasing trend for TSPAN3(Y252A), suggesting that these residues contribute to TSPAN3-dependent accumulation or remodeling of LC3-positive compartments.

We next asked whether T75 and Y252 also contribute to the incorporation of TSPAN3 into secretory MVBs. TIRF microscopy revealed a strong reduction in MVB–plasma membrane fusion events for both TSPAN3(T75A) and TSPAN3(Y252A) compared with TSPAN3 WT (Figure 4K). Because both mutants retained a predominantly intracellular vesicular localization (Figure 4E), the reduction in fusion was unlikely to result solely from gross mislocalization. These findings suggest that T75 and Y252 contribute to the formation, trafficking, or secretory competence of TSPAN3-positive MVBs.

Having established a functional association between TSPAN3 and autophagy-related machinery, we next investigated how perturbation of the autophagy–endolysosomal pathway affects TSPAN3-positive EV secretion. We inhibited VPS34 using SAR405 and quantified EV secretion using the Nluc-based assay (Figure 4L). Unexpectedly, rather than reducing secretion, SAR405 increased the release of both CD63-and TSPAN3-positive EVs, with a more pronounced increase observed for TSPAN3. Moreover, SAR405 further enhanced TSPAN3-positive EV secretion in the presence of BafA1.

We next examined the corresponding cellular Nluc levels to determine whether the increased extracellular signal could be explained by changes in cellular reporter abundance (Figure 4M). BafA1 increased the cellular levels of both CD63 and TSPAN3. SAR405 also increased cellular CD63 levels, whereas the cellular TSPAN3 pool remained largely unchanged despite the pronounced increase in TSPAN3-positive EV secretion. Combined SAR405 and BafA1 treatment did not produce a comparable additive increase in cellular TSPAN3 levels. Thus, the enhanced release of TSPAN3 following VPS34 inhibition cannot be explained by a corresponding increase in cellular TSPAN3 abundance, consistent with an altered distribution of TSPAN3 between intracellular and secretory fates.

To further distinguish the contribution of the ATG8 conjugation machinery from that of VPS34-dependent membrane trafficking, we next inhibited ATG7 using ATG7-IN-2. In contrast to VPS34 inhibition, ATG7-IN-2 caused a small but significant reduction in both CD63-and TSPAN3-positive EV secretion (Figure 4N), suggesting that ATG7-dependent machinery contributes to efficient EV release but is not absolutely required for secretion.

Analysis of the corresponding cellular Nluc levels revealed a differential effect on CD63 and TSPAN3. ATG7-IN-2 reduced the cellular CD63 pool, whereas cellular TSPAN3 levels were not significantly altered by ATG7 inhibition alone. Strikingly, however, ATG7-IN-2 completely suppressed the BafA1-induced increase in cellular CD63 and TSPAN3 signal. Thus, while basal cellular TSPAN3 abundance was relatively insensitive to ATG7 inhibition, the accumulation of TSPAN3 induced by lysosomal inhibition required intact ATG7-dependent machinery. Together, these results suggest that ATG7 activity makes a modest contribution to basal CD63-and TSPAN3-positive EV secretion. More strikingly, ATG7 inhibition prevented the cellular accumulation of both tetraspanins following lysosomal inhibition, indicating that the BafA1-induced expansion of these intracellular pools requires ATG7-dependent machinery.

Collectively, these findings identify TSPAN3 as a molecular link between secretory MVBs and autophagy-related membrane trafficking. TSPAN3 associates with LC3-positive compartments and a selective ATG8-related protein network, while T75 and Y252 are required for both this association and efficient delivery to fusion-competent secretory MVBs. The differential effects of VPS34 and ATG7 inhibition further demonstrate that autophagy-related pathways regulate the balance between intracellular retention and secretion of TSPAN3-positive compartments.

**Figure 3:**
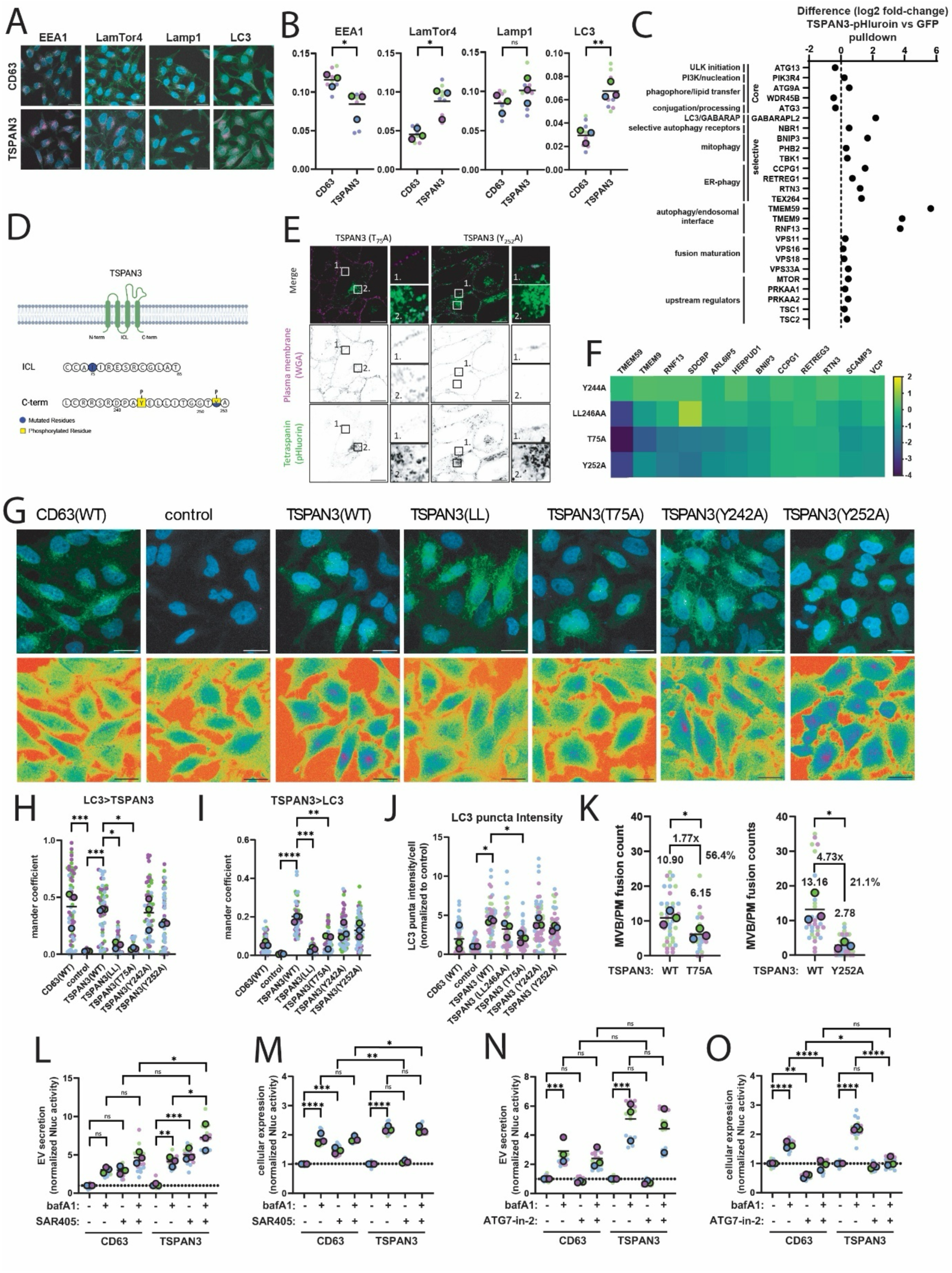
TSPAN3 engages autophagy-related machinery during trafficking to secretory MVBs. (A) Representative fluorescence images of CD63-HALO and TSPAN3-HALO co-stained with markers of distinct intracellular compartments: EEA1 (early endosomes), LAMTOR4 (late endosomal/lysosomal compartments), LAMP1 (lysosomes), and LC3 (autophagy-related membranes). (B) Quantification of the colocalization of CD63-HALO and TSPAN3-HALO with the compartment markers shown in (A). (C) Bargraph showing autophagy/endosomal related proteins of TSPAN3(WT)-pHluorin quantified using GFP affinity purification–mass spectrometry approach. (D) Schematic representation of TSPAN3 topology and the predicted LC3-interacting region (LIR) motif(s), including the residues targeted for mutagenesis. (E) Representative super-resolution fluorescence images of TSPAN3 WT, TSPAN3(T75A), and TSPAN3(Y252A), co-stained with wheat germ agglutinin (WGA) to visualize the plasma membrane. (F) Heatmap showing the effect of TSPAN3(Y243A), TSPAN3(LL246AA), TSPAN3(T75A), and TSPAN3(Y252A), compared to TSPAN3(WT) on autophagy/endosomal related proteins quantified using GFP affinity purification-mass spectrometry approach using TSPAN-pHluorin fusion proteins. (G) Representative fluorescence images of CD63 WT and TSPAN3 WT, TSPAN3(T75A), and TSPAN3(Y252A) co-stained for LC3. (H, I) Quantification of LC3 colocalization with CD63 WT and TSPAN3 WT, TSPAN3(T75A), and TSPAN3(Y252A) from the images represented in (G). (J) Quantification of cellular LC3 signal in cells expressing CD63 WT or TSPAN3 WT, TSPAN3(T75A), and TSPAN3(Y252A). (K) Quantification of MVB/plasma membrane fusion events for TSPAN3 WT, TSPAN3(T75A), and TSPAN3(Y252A) using pHluorin-based TIRF microscopy. (L) Nluc-based quantification of EV secretion from HA-Nluc-CD63 and HA-Nluc-TSPAN3 cells following treatment with Bafilomycin A1 (BafA1), SAR405, or the combination of BafA1 and SAR405. (M) Corresponding cellular Nluc levels for the conditions shown in (L). (N) Nluc-based quantification of EV secretion from HA-Nluc-CD63 and HA-Nluc-TSPAN3 cells following treatment with BafA1, ATG7-IN-2, or the combination of BafA1 and ATG7-IN-2. (O) Corresponding cellular Nluc levels for the conditions shown in (M). Data are from n = 3 independent experiments. Each big point represents the mean of an individual experiment. Each small point represents an individual quantification. Statistical significance was determined using one-way anova; ns, not significant; *P < 0.01, *P* < 0.001, *P < 0.0001. Scale bars, 20 µm unless otherwise indicated.

### CD63-and TSPAN3-associated EVs display distinct cargo signatures consistent with differential membrane origins

Having established that CD63 and TSPAN3 reach secretory MVBs through distinct intracellular trafficking routes, we next asked whether these differences are reflected in the molecular composition of the EVs they generate. To address this, we analyzed EV cargo biotinylated in proximity to uID-CD63 or uID-TSPAN3 prior to secretion (Figure 5A). Cells expressing uID-CD63 or uID-TSPAN3 were incubated with biotin for 24 h, after which EVs were isolated from conditioned medium by size-exclusion chromatography. Biotinylated EV proteins were enriched by Neutravidin affinity purification and analyzed by quantitative mass spectrometry. EVs derived from wild-type cells were processed in parallel to define proteins specifically enriched in the uID-CD63-and uID-TSPAN3-associated EV cargo.

Comparison with wild-type EVs identified extensive enrichment of biotinylated proteins in both uID-CD63 and uID-TSPAN3 EVs, indicating that proteins labeled in proximity to either tetraspanin were efficiently incorporated into secreted EVs (Figure 5B,C). Although a substantial fraction of the enriched proteins was shared between the two conditions, the relative enrichment profiles differed markedly, suggesting that CD63-and TSPAN3-associated EVs acquire cargo from distinct membrane environments during biogenesis. We therefore next asked whether these differences could be resolved at the level of the cellular compartments represented within the EV cargo.

Gene set enrichment analysis of GO Cellular Component terms revealed a pronounced plasma membrane signature in CD63-associated EV cargo (Figure 5D). Terms including lateral plasma membrane, cell surface, basolateral plasma membrane, and external side of plasma membrane were among the most strongly enriched categories in CD63 EVs. In contrast, endosomal and endocytic-vesicle-associated terms were relatively more prominent in TSPAN3 EV cargo. In particular, endocytic vesicle membrane was significantly enriched in TSPAN3-associated EVs, whereas the corresponding enrichment was substantially weaker in CD63 EVs. Similarly, terms associated with the recycling endosome, endosome membrane, and late endosome consistently showed higher enrichment scores in the TSPAN3 dataset than in the CD63 dataset. Although plasma membrane-associated terms were also enriched in the TSPAN3 dataset, their normalized enrichment scores were consistently lower than those observed for CD63. These findings indicate that CD63-associated EV cargo retains a stronger cell-surface/plasma-membrane signature, whereas TSPAN3-associated EV cargo shows a comparatively greater contribution from endocytic and endosomal compartments.

We next asked whether individual EV cargo proteins similarly distinguished CD63-and TSPAN3-associated EVs (Figure 5E). CD63-selective cargo included several plasma membrane, cell-surface, and membrane-associated proteins, including SLC29A2, STRA6, NPTN, CLMP, SLC5A3, F11R, and HLA-C. In contrast, TSPAN3-selective cargo included proteins associated with endosomal and endolysosomal trafficking, including LAMP1, RAB8B, MFSD8, and RRAGC, together with additional proteins preferentially enriched in TSPAN3-associated EVs. Thus, the identities of the selectively enriched proteins recapitulated the compartment-level differences identified by GSEA.

Together, these findings indicate that the distinct intracellular trafficking routes followed by CD63 and TSPAN3 are reflected in the molecular composition of the EVs they generate. The strong plasma membrane signature of CD63-associated EV cargo is consistent with extensive trafficking through the cell surface, potentially allowing both direct plasma membrane budding and subsequent internalization into the endosomal system. In contrast, the comparatively stronger endosomal and endocytic signature of TSPAN3-associated EV cargo supports its origin from a more intracellular secretory pathway. Thus, the EV proteome retains a molecular signature of the distinct trafficking histories followed by CD63 and TSPAN3, providing independent support for the differential membrane origins of these EV populations.

**Figure 4:**
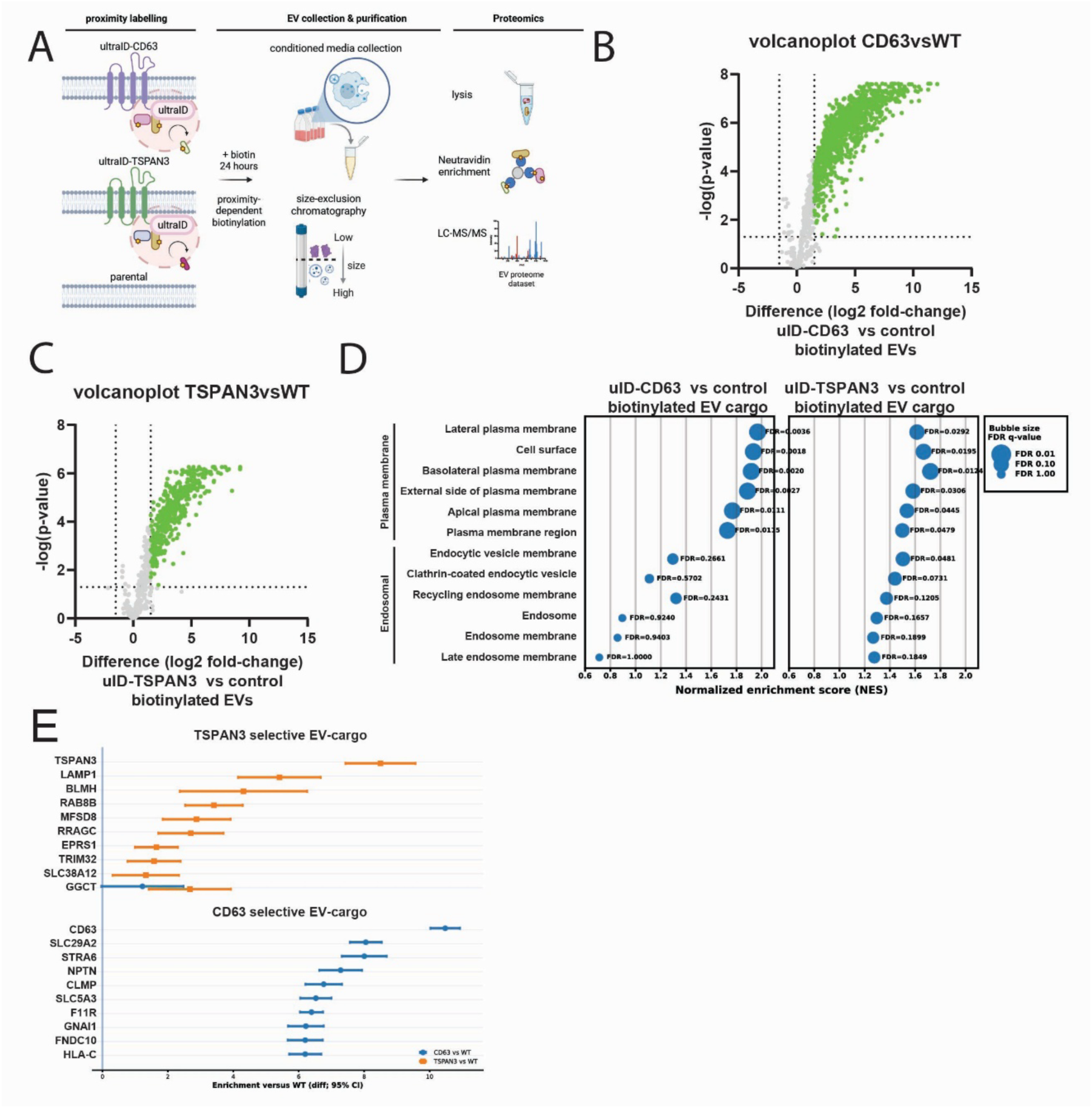
CD63-and TSPAN3-associated EVs contain distinct cargo signatures consistent with differential membrane origins. (A) Schematic overview of the biotinylated EV-cargo proteomics workflow. Cells expressing uID-CD63 or uID-TSPAN3 were incubated with biotin for 24 h, after which extracellular vesicles (EVs) were isolated from conditioned medium by size-exclusion chromatography. Biotinylated EV proteins were enriched by Neutravidin affinity purification and analyzed by quantitative mass spectrometry. EVs from wild-type (WT) cells were processed in parallel as a control. (B,C) Volcano plots showing differential protein abundance in biotinylated EV cargo from uID-CD63 versus WT (B) and uID-TSPAN3 versus WT (C). (D) Gene set enrichment analysis (GSEA) of GO Cellular Component terms associated with the plasma membrane and endosomal compartments. Normalized enrichment scores (NES) are shown for CD63-and TSPAN3-associated EV cargo, with bubble size representing the FDR q-value. CD63-associated EV cargo showed stronger enrichment of plasma membrane and cell-surface terms, whereas endosomal and endocytic-vesicle terms were relatively more prominent in TSPAN3-associated EV cargo. (E) Top 10 condition-selective proteins enriched in CD63-or TSPAN3-associated EV cargo. Points indicate differential enrichment relative to WT and whiskers indicate 95% confidence intervals. Proteins were selected based on significant enrichment in the indicated EV population and preferential enrichment relative to the alternative condition. Together, these analyses reveal distinct EV-cargo profiles associated with CD63 and TSPAN3, consistent with differential contributions of plasma membrane and endosomal trafficking routes to EV biogenesis.

## Discussion

### TSPAN3 reveals heterogeneity among secretory MVBs and MVB-derived EVs

Our findings identify TSPAN3 as a marker of a secretory MVB population that is molecularly and functionally distinct from canonical CD63-positive compartments, highlighting substantial heterogeneity within the endosomal EV pathway. Increasing evidence indicates that molecular heterogeneity is established already within the late endosomal/lysosomal system. Multiplexed single-organelle imaging has revealed distinct late endosome/lysosome populations defined by combinations of membrane proteins, including CD63 and LAMTOR4, while functionally distinct secretory MVB populations have been distinguished by their endosomal origin, maturation state and molecular machinery^22,23^. This diversity includes Rab11a-positive recycling endosomal compartments that give rise to a distinct EV population^24^, as well as a Rab27-positive pre-endolysosomal subset of CD63-positive MVBs selectively competent for secretion^8^. More recently, phosphoinositide remodeling^25^ and differential engagement of Bro1-family proteins^26^ have been implicated in directing individual MVBs toward either plasma-membrane fusion or lysosomal degradation. Together, these studies support the existence of multiple secretory MVB populations with distinct molecular identities and cellular fates. Our findings extend this framework by identifying tetraspanin composition as an additional feature that distinguishes secretory MVB identity (Figure 1).

Previous work using EV-engineering approaches suggested that TSPAN3 marks an EV population with low abundance of the canonical tetraspanins CD9, CD63 and CD81 and a proteomic composition distinct from CD63-engineered EVs^27^. Our study provides the intracellular context for these observations by showing that TSPAN3 localizes to a secretory MVB population that is largely distinct from canonical CD63-positive compartments and reaches these compartments through a different trafficking route. Consistent with this, biotinylated EV-cargo proteomics revealed distinct molecular signatures in CD63-and TSPAN3-associated EVs supporting the idea that these tetraspanins are associated with different EV populations. Despite remaining comparatively understudied in EV biology, TSPAN3 is a conserved vertebrate tetraspanin and was recently included among the top 100 abundant proteins proposed as a reference marker panel for small EVs^5^. Taken together, our findings indicate that TSPAN3 is not simply an additional EV marker but identifies a previously underappreciated population of secretory MVBs that contributes to EV heterogeneity.

### Trafficking history may determine EV identity

The distinct intracellular trafficking routes followed by CD63 and TSPAN3 raise the possibility that the membrane environments encountered before secretion contribute to the molecular composition of the EVs they generate. Although our study does not establish a causal relationship between trafficking route and EV composition, the correspondence between intracellular localization and EV cargo is consistent with such a model.

CD63 undergoes dynamic trafficking between the plasma membrane and endolysosomal system, with its C-terminal YXXΦ motif mediating adaptor-complex-dependent internalization and lysosomal targeting^6,7^. Consistent with previous work, we observed rapid internalization of surface CD63 and a strong dependence of secretory MVB fusion on its YXXΦ motif. By contrast, although TSPAN3 also contains overlapping YXXΦ and dileucine motifs, mutation of the YXXΦ tyrosine had only a modest effect, whereas disruption of the LL-containing region strongly impaired trafficking to secretory MVBs. Because mutation of the tyrosine alone is generally sufficient to abolish canonical AP-complex recognition, these findings argue that classical YXXΦ-dependent trafficking contributes only modestly to TSPAN3 sorting, whereas determinants within the overlapping LL-containing region play the dominant role. Together, these findings indicate that CD63 and TSPAN3 reach secretory MVBs through distinct sorting mechanisms.

These differences were reflected in the molecular composition of the secreted EV populations. CD63-associated EVs were enriched for plasma membrane and cell-surface proteins, whereas TSPAN3-associated EVs showed comparatively stronger enrichment of endosomal and endolysosomal proteins, both at the level of GO terms and individual cargo molecules. Together, these observations suggest that EV proteomes retain molecular signatures of the membrane nanodomains associated with individual tetraspanins and of the intracellular trafficking routes followed by those domains. More generally, our findings support the emerging view that tetraspanins actively organize membrane trafficking rather than merely serving as EV markers^29^.

An important limitation is that these experiments do not directly establish the biogenesis route of individual extracellular TSPAN3-positive vesicles. Although the endosomal signature of TSPAN3-associated EV cargo, together with its localization to ILV-containing MVBs and direct visualization of TSPAN3-positive MVB fusion events, supports a substantial contribution from the endosomal pathway, EV cargo composition alone cannot distinguish vesicles released by MVB fusion from those budding directly from the plasma membrane. A contribution from plasma membrane-derived TSPAN3-positive EVs therefore cannot be excluded.

### TSPAN3 engages a selective autophagy-related membrane network that influences secretory MVB fate

The distinct intracellular trafficking of TSPAN3 was accompanied by an unexpected association with autophagy-related machinery. TSPAN3 showed substantially greater association with LC3-positive compartments than CD63, while its associated proteome contained a selective group of ATG8-, selective-autophagy-, and endosomal trafficking proteins, including GABARAPL2, NBR1, BNIP3, CCPG1, TMEM59, TMEM9 and RNF13. Importantly, this signature did not represent broad enrichment of the core canonical autophagy machinery, suggesting that TSPAN3 may intersect with a specialized autophagy-related membrane pathway rather than simply localizing to conventional autophagosomes. Mutation of T75 or Y252 altered the association of TSPAN3 with several proteins involved in autophagy-related and endosomal trafficking, indicating that these residues contribute to the molecular organization of the TSPAN3-associated compartment. Consistent with this, both T75A and Y252A reduced the association of TSPAN3 with LC3-positive compartments and strongly impaired TSPAN3-positive MVB fusion. Together, these observations identify a functional link between TSPAN3 trafficking and a selective LC3/ATG8-associated membrane network, although the molecular basis of this relationship remains unresolved.

An interesting possibility is that LC3/ATG8 proteins are recruited directly to TSPAN3-positive endosomal membranes rather than TSPAN3 entering the canonical double-membrane autophagosome pathway. Autophagy proteins increasingly have recognized functions on single-membrane endolysosomal compartments through non-canonical ATG8 conjugation pathways, including conjugation of ATG8 to single membranes (CASM)^30^ and LC3-associated endocytosis^31^. In these pathways, components of the LC3-conjugation machinery are recruited to endosomal or lysosomal membranes without requiring formation of a canonical autophagosome. A particularly relevant precedent is TMEM59, which recruits ATG16L1 and promotes LC3 labeling of its own endosomal compartment^32^. Notably, TMEM59 was also strongly enriched in the TSPAN3-associated proteome. The LC3-conjugation machinery has furthermore been implicated directly in EV biology through LC3-dependent EV loading and secretion (LDELS), in which ATG7-dependent LC3 conjugation regulates the incorporation of selected cytosolic and transmembrane cargoes into EVs^33,34^. Collectively, these studies provide a plausible framework in which TSPAN3-positive compartments engage selected components of the ATG8 machinery without necessarily entering the canonical autophagy pathway, consistent with the effects observed upon VPS34 inhibition. However, whether this indeed represents non-canonical ATG8 conjugation directly on TSPAN3-positive endosomes, convergence with an autophagosomal compartment, or communication between otherwise distinct membrane pathways remains to be determined.

The pharmacological perturbations further indicate that different components of the autophagy/endolysosomal system regulate distinct aspects of TSPAN3 trafficking. Inhibition of lysosomal acidification with BafA1 strongly enhanced TSPAN3-positive EV secretion, consistent with the idea that disruption of lysosomal degradation can redirect intracellular membrane compartments toward secretion. Secretory competence has been proposed to emerge during MVB maturation, with CD63-positive secretory MVBs representing a non-proteolytic pre-endolysosomal population that undergoes a Rab7a–Arl8b–Rab27a GTPase transition before plasma membrane fusion^8^. More direct evidence for competition between degradative and secretory MVB fates comes from disruption of the BORC–ARL8– HOPS machinery required for MVB–lysosome fusion, which increases the number of ILV-containing MVBs, promotes their fusion with the plasma membrane, and enhances exosome secretion^9^. More recently, differential engagement of the Bro1-family proteins ALIX and PTPN23 was similarly shown to influence whether MVBs undergo secretion or lysosomal degradation^26^. Together, these studies support the concept that TSPAN3-positive compartments occupy a trafficking branch point at which the balance between lysosomal degradation and plasma membrane fusion determines their secretory output.

The response to VPS34 inhibition further supports a role for endolysosomal compartment maturation in regulating TSPAN3-positive EV secretion. Rather than reducing secretion, SAR405 markedly increased the release of TSPAN3-positive EVs, including in the presence of BafA1, while cellular TSPAN3 abundance remained largely unchanged. VPS34 generates PI(3)P required for both autophagosome biogenesis and endolysosomal trafficking, and SAR405 has been shown to disrupt trafficking from late endosomes to lysosomes in addition to inhibiting autophagy^35^. The enhanced secretion observed following VPS34 inhibition therefore likely reflects broader perturbation of endolysosomal maturation rather than inhibition of autophagy alone.

By contrast, inhibition of ATG7 produced only a modest reduction in EV secretion while preventing the BafA1-induced accumulation of intracellular CD63 and TSPAN3. These findings indicate that ATG7-dependent LC3 conjugation contributes to the homeostasis of tetraspanin-positive compartments but is unlikely to be the principal determinant of their secretory output. Instead, the divergent effects of VPS34 and ATG7 inhibition suggest that distinct components of the autophagy/endolysosomal network regulate different aspects of TSPAN3-positive compartment biology.

Taken together, our data argue against a simple linear relationship between autophagy and EV secretion. Rather, our findings support a model in which TSPAN3 marks a distinct endosomal membrane population that selectively associates with LC3/ATG8-related machinery while remaining subject to broader pathways governing endolysosomal maturation and the balance between degradative and secretory MVB fates. Whether LC3/ATG8-associated proteins directly regulate the formation or function of these compartments, or instead reflect communication between parallel membrane-trafficking pathways, remains an important question for future studies.

### Concluding model

Together, our findings support a model in which TSPAN3 identifies a population of secretory MVBs that is distinct from canonical CD63-positive compartments. Compared with CD63, TSPAN3 reaches these compartments through a different intracellular trafficking route and associates with a selective LC3/ATG8-related molecular network, while generating EVs with a distinct molecular composition. Although the relationship between these features remains to be established, their convergence identifies TSPAN3-positive MVBs as a molecularly distinct branch of the secretory endosomal system. TSPAN3 therefore provides a molecular handle to resolve secretory MVB heterogeneity and investigate how distinct intracellular membrane compartments contribute to extracellular vesicle diversity.

## Supplementary Figures

**Supplementary Figure 1.**
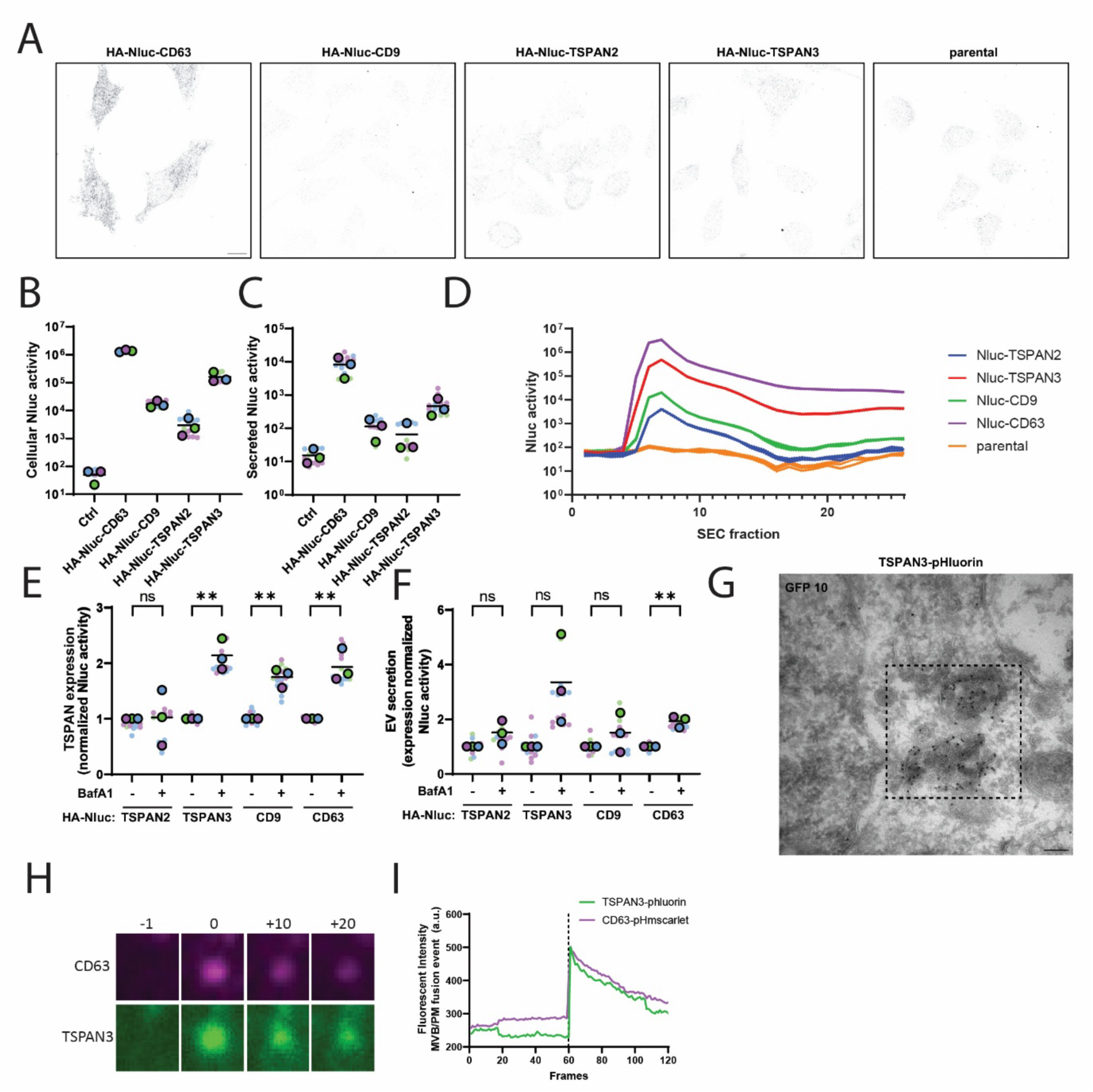
Characterization of HA-Nluc–tetraspanin reporters and TSPAN3-positive secretory events. (A) Representative immunofluorescence images of cells expressing HA-Nluc-CD63, HA-Nluc-CDG, HA-Nluc-TSPAN2, or HA-Nluc-TSPAN3, stained for the HA tag; parental cells are shown as a negative control. (B) Cellular NanoLuc (Nluc) activity measured in cells expressing the indicated HA-Nluc– tetraspanin constructs. (C) Secreted Nluc activity detected in conditioned medium from cells expressing the indicated constructs. (D) Size-exclusion chromatography (SEC) fractionation of conditioned medium, showing the distribution of secreted Nluc activity across SEC fractions for HA-Nluc-TSPAN2, HA-Nluc-TSPAN3, HA-Nluc-CD9, HA-Nluc-CD63, and parental cells. (E) Effect of bafilomycin A1 (BafA1) treatment on cellular Nluc activity of the indicated HA-Nluc–tetraspanin constructs, corresponding to Figure 1B. Values are normalized to the respective untreated condition. (F) Effect of BafA1 treatment on EV-associated Nluc secretion normalized to cellular Nluc activity, corresponding to Figure 1B. (G) Representative electron micrograph showing immunogold labeling of TSPAN3-pHluorin, corresponding to Figure 1E. The boxed region indicates a TSPAN3-positive multivesicular body. (H) Representative time series of a CD63-pHmScarlet/TSPAN3-pHluorin double-positive MVB–plasma membrane fusion event, with time shown relative to the onset of fusion, corresponding to Figure 1G. (I) Corresponding fluorescence-intensity traces of CD63-pHmScarlet and TSPAN3-pHluorin during the fusion event shown in (H); the dashed line indicates the onset of fusion. Data points in (B), (C), (E), and (F) represent individual biological replicates; horizontal lines indicate the mean. ns, not significant; **P < 0.01.

**Supplementary Figure 2:**
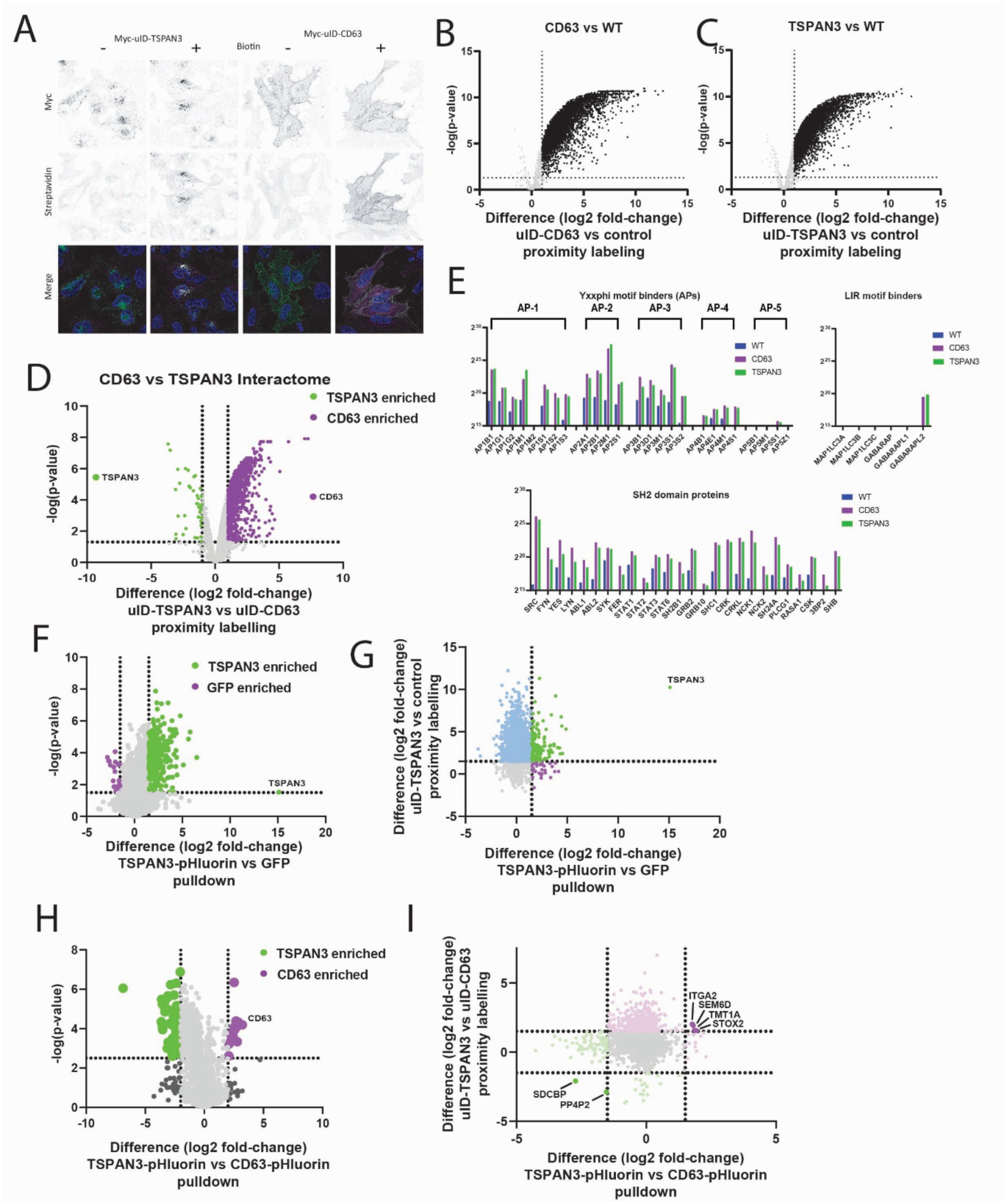
Proteomic characterization of the TSPAN3 and CD63 molecular environments and comparison with affinity-purification proteomics. (A) Representative fluorescence images of cells expressing Myc-ultraID-TSPAN3 or Myc-ultraID-CD63 in the absence or presence of biotin. Myc-ultraID fusion proteins were detected by anti-Myc staining (green), biotinylated proteins were visualized using fluorescent streptavidin (magenta), and nuclei are shown in blue. (B–C) Volcano plots showing differential protein enrichment in the proximity proteomes of Myc-ultraID-CD63 versus wild-type cells (B) and Myc-ultraID-TSPAN3 versus wild-type cells (C). (D) Volcano plot comparing the Myc-ultraID-TSPAN3 and Myc-ultraID-CD63 proximity proteomes, corresponding to the comparison shown in Figure 3C. Proteins enriched in the TSPAN3 and CD63 proximity proteomes are indicated in green and purple, respectively. (E) Relative abundance of selected trafficking-and interaction-related protein families detected in wild-type, Myc-ultraID-CD63, and Myc-ultraID-TSPAN3 proteomes, including components of the AP-1 to AP-5 adaptor protein complexes, LIR motif-binding ATG8-family proteins, and SH2 domain-containing proteins. (F) Volcano plot comparing proteins isolated by TSPAN3-pHluorin affinity purification with the GFP control. Proteins enriched in the TSPAN3-pHluorin pulldown are indicated in green and proteins enriched in the GFP control in purple. (G) Comparison of protein enrichment in the TSPAN3-pHluorin versus GFP pulldown and Myc-ultraID-TSPAN3 versus control proximity-labeling datasets. Each point represents an individual protein and is positioned according to its log2 fold-change in the two independent proteomic approaches. (H) Volcano plot comparing the TSPAN3-pHluorin and CD63-pHluorin affinity-purification proteomes. Proteins preferentially enriched with TSPAN3-pHluorin or CD63-pHluorin are indicated in green and purple, respectively. (I) Comparison of differential protein enrichment between TSPAN3 and CD63 obtained by affinity-purification and proximity-labeling proteomics. Each point represents an individual protein and is positioned according to its log2 fold-change in the two proteomic approaches; selected proteins showing concordant or discordant enrichment are indicated. Dotted lines indicate the significance and fold-change thresholds used for differential enrichment.

**Supplementary Figure 3:**
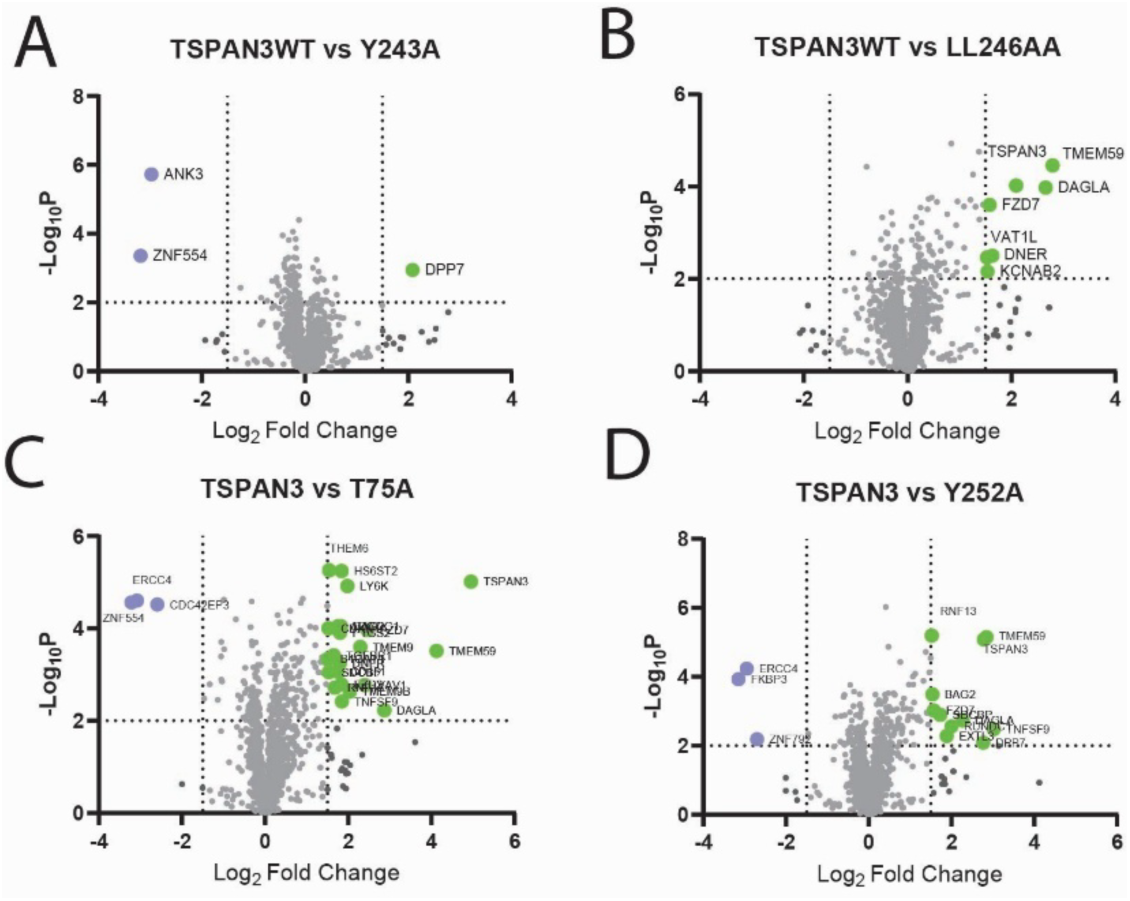
Proteomic comparison of wild-type TSPAN3 and trafficking-motif mutants. (A–D) Volcano plots showing differential protein enrichment between wild-type TSPAN3 (TSPAN3 WT) and the indicated TSPAN3 mutants: Y243A (A), LL246AA (B), T75A (C), and Y252A (D). Positive log2 fold-change values indicate proteins enriched with TSPAN3 WT relative to the respective mutant, whereas negative values indicate preferential enrichment with the mutant. Selected differentially enriched proteins are highlighted and labeled. Dotted lines indicate the fold-change and significance thresholds used to define differential enrichment.

## Notes

### Competing Interest Statement

The authors have declared no competing interest.

### Funding

- Dutch Research Council: VI.Vidi.213.106
- European Research Council: 101075975

